# ProStab: Prediction of protein stability change upon mutations by protein language and inverse folding models

**DOI:** 10.1101/2025.08.11.669595

**Authors:** Hong Tan, Xiaowei Wei, Shenggeng Lin, Xueying Mao, Junwei Chen, Heqi Sun, Yufang Zhang, Zhenghong Zhou, Dong-Qing Wei, Shuangjun Lin, Yi Xiong

## Abstract

Predicting protein stability change upon mutation is critical for protein engineering, yet remains limited by the modeling assumptions of physics-based methods and the generalization bottlenecks of data-driven approaches. We present ProStab, a deep learning framework that integrates sequence- and structure-based information, including the mutation-aware sequence embeddings from protein language models and the geometric features extracted via an inverse folding model. Trained on the large-scale Megascale dataset, ProStab demonstrates strong performance across diverse test sets and robust generalization across distribution shifts between the training and test sets. In head-to-head comparisons, ProStab outperforms all state-of-the-art methods with consistently higher Spearman correlation and precision. To evaluate its practical utility, we experimentally validated ProStab-predicted mutations on the model enzyme transaminase. Among the 16 successfully expressed variants, 4 exhibited improved thermal stability. Remarkably, the 1st top-ranked predicted mutation yielded the highest observed enzymatic activity, retaining three-fold that of the wild type after 10 minutes at 40 °C. To facilitate broader application, a publicly accessible web server has been developed. We envisage that ProStab provides a scalable and accurate platform for intelligent protein stability design.

## Introduction

Thermal stability is a fundamental property of proteins that determines their functionality and lifespan in both natural and engineered environments^1^. However, many wild-type proteins exhibit only marginal stability under physiological or industrial conditions^2^, limiting their utility in applications such as biocatalysis^3,4^, biotherapeutics^5^, and biosensing^6^. Enhancing thermostability has therefore become a key focus in protein engineering. Thermal stability is typically quantified by the Gibbs free energy difference (ΔG) between the folded and unfolded states, while the effect of mutations is assessed as the change in stability (ΔΔG) between the wide-type and mutation states. Accurately predicting how sequence variations determine ΔΔG is essential for the rational design of thermostable protein variants in biomedical and industrial contexts.

Experimental approaches such as directed evolution^7^ and mutational scanning^8^ offer effective methods for identifying stabilizing mutations. However, their scalability is constrained by cost, labor, and throughput^9,10^. Thermostabilizing mutations are rare even at the single-substitution level, rendering exhaustive exploration impractical. As a result, accurate and scalable computational prediction of ΔΔG has become an indispensable component of a modern protein engineering workflow.

Among computational strategies for ΔΔG prediction, physics-based methods such as FoldX^11^ and Rosetta^12^ remain widely adopted for their mechanistic interpretability and demonstrated effectiveness across diverse protein families. However, their reliance on fixed-backbone models and discrete side-chain rotamer substitutions limits their ability to capture mutation-induced local structural adaptations and broader conformational rearrangements. Moreover, their high computational cost poses a significant barrier to large-scale saturation mutagenesis, rendering them impractical for high-throughput variant screening.

Recent advances in deep learning (DL) have enabled new strategies for ΔΔG prediction, broadly categorized into sequence-based^13–16^, structure-based^17–20^, and sequence-structure hybrid models^6,21^. Sequence-based models, such as PROSTATA^13^, leverage protein language model embeddings, effectively capture evolutionary constraints but lack direct grasp in the physicochemical principles that govern protein stability. Structure-based models, exemplified by ThermoNet^17^, attempt to infer mutational effects by comparing mutant and wild-type conformations. However, they often underestimate structural perturbations caused by single-point mutations. Hybrid models, such as ThermoMPNN^6^ and SPURS^21^, integrate sequence and structural information but do not explicitly model the mutated sequence or structure, limiting their capacity to capture mutation-induced conformational rearrangements.

In addition to limitations in model architecture and representation, the quality and diversity of training data also pose significant challenges for ΔΔG prediction. Most existing stability datasets are biased towards alanine scanning experiments, in which various types of amino acids are mutated to alanine, with limited representation of reverse or non-alanine substitutions^18^. Moreover, many deep learning models are trained on small datasets, such as S2648, which contains only 2,648 training examples and S669 for independent testing, substantially limiting their ability to generalize^22^. Moreover, most models lack web-lab experimental validation, raising concerns about their reliability and practical utility in real-world protein design settings.

To address these challenges, we developed ProStab, a deep learning framework that integrates sequence- and structure-derived features for accurate prediction of ΔΔG for protein point mutations given an initial structure. ProStab combines representations from a protein language model^23^ applied to both wild-type and mutant sequences, and from the inverse folding model ProteinMPNN^24^ applied to the wild-type structure. It jointly models two sources of information: mutation-specific effects, captured as embedding differences at the substitution site between wild-type and mutant sequences; and site-specific priors, derived from the wild-type sequence and structure, which reflect the local context and substitutional tolerance. By integrating these complementary signals, ProStab predicts the stability impact of single-residue mutations without requiring explicit structural modeling of the mutant. Trained on the largest available dataset (the Megascale dataset), which consists mainly of small, single-domain proteins, ProStab provides a scalable and generalizable solution for mutation effect prediction, performing robustly on longer, structurally diverse proteins. To validate the effect of mutations on protein stability predicted by ProStab, we carry out wet experiments to investigate the actual effect of the top-ranked predicted mutations of amine transaminase (ATA) from *Exophiala xenobiotica* (Ex). ATAs are suitable for the synthesis of industrial relevant chiral amines^25–28^, due to their substrates including prochiral ketones without α-carboxylic acid group^29,30^. Remarkably, the ATA from *Ex* (Ex-ATA) has shown excellent catalytic efficiency and high R-enantiopreference^31^. However, the thermostability of Ex-ATA needs to be further improved for industrial application. We assessed 20 top-ranked predicted mutations of Ex-ATA (PDB ID: 6FTE). Among the 16 variants that were successfully expressed, four exhibited enhanced thermal stability. Notably, the 1st top-ranked variant retained over threefold higher enzymatic activity than the wild type after 10 minutes at 40 °C. In sum, this case exemplifies how ProStab-predicted mutations can enhance thermal stability, thereby preserving enzymatic activity under elevated temperatures.

To support broader use, we have made ProStab freely available via a publicly accessible web server at https://ails.sjtu.edu.cn/prostab, enabling rapid, high-throughput ΔΔG prediction and facilitating scalable protein engineering in both biomedical and industrial contexts.

## Results

### Overview of ProStab

The overall architecture of ProStab is illustrated in Fig. 1b. ProStab employs an end-to-end framework that seamlessly integrates both sequence- and structural-derived features to predict ΔΔG. It takes three types of information as the input: the wild-type protein sequence, the corresponding mutant sequence, and the three-dimensional structure of the wild-type protein.

**Figure 1.**
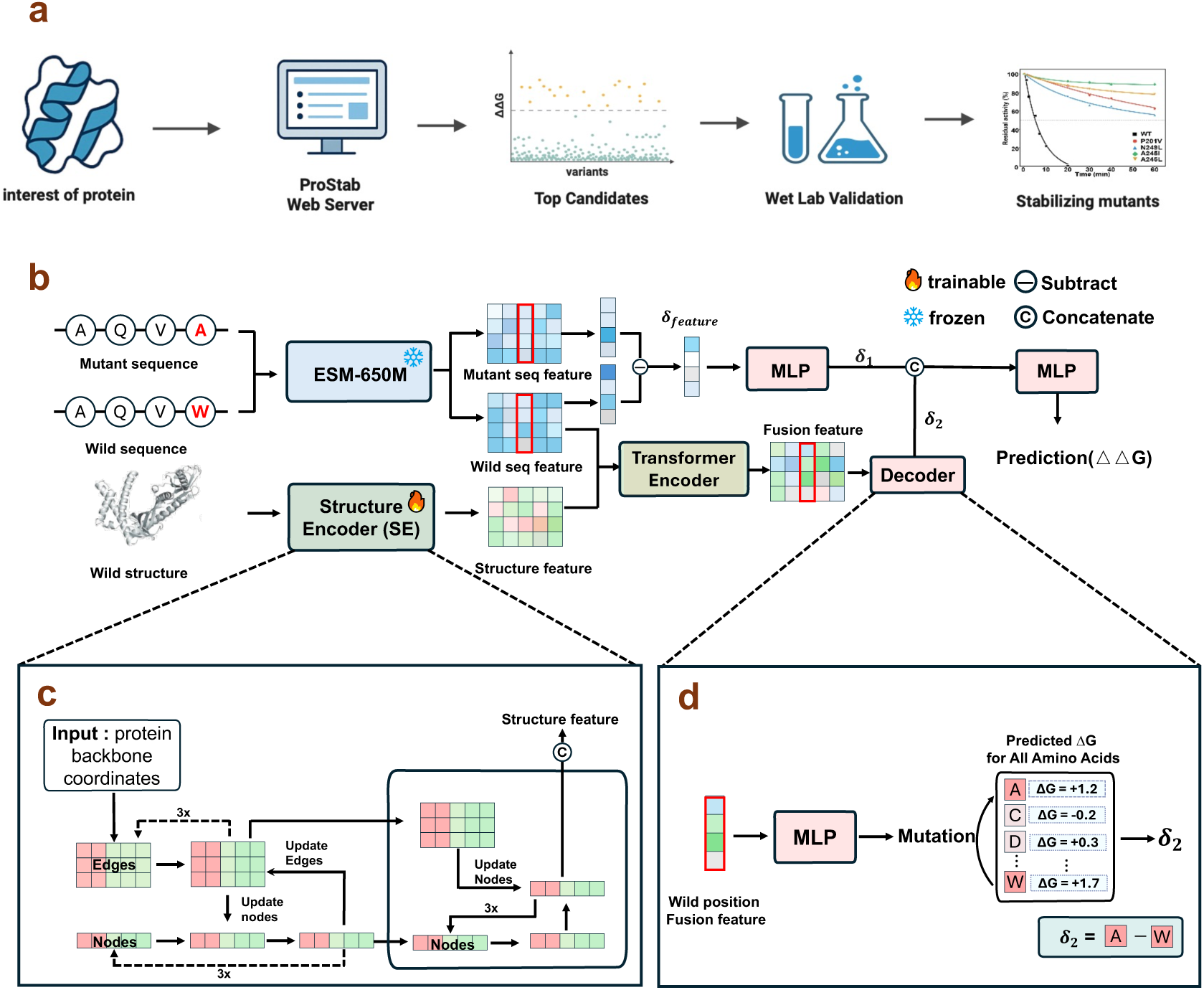
Overview of ProStab. (a) The Pipeline of protein stabilization via computational and experimental screening. (b) Overview of the ProStab framework. The model integrates two sources of information for ΔΔG prediction: mutation-specific effects, derived from embedding differences between wild-type and mutant sequences; and site-specific priors, learned from the wild-type sequence and structure. (c) Backbone coordinates of the input protein are processed by the Structure Encoder to generate residue-level structural representations. (d) Architecture of the decoder module. It learns site-specific priors from wild-type context and uses the fusion feature at the substitution site to estimate the local ΔΔG effect.

For sequence representation, we utilize ESM2-650M^23^ to extract amino acid-level features from both the wild-type and mutant sequences. The token representations at the mutation site are extracted, and their difference (denoted as δ_feature_) is computed to capture the localized embedding-level change caused by the mutation.

To extract structural information, we implement a Structure Encoder (SE) adapted from ProteinMPNN^24^, which derives geometric features from the backbone coordinates of the wild-type protein.

The wild-type sequence features and structural features are then integrated via a Transformer Encoder^32^ to generate a joint representation that captures the contextual relationship between sequence and structure. This representation is subsequently processed by a specialized decoder to assess the positional impact of substituting the wild-type amino acid with the mutant residue, yielding a prediction denoted as δ_2_. In parallel, δ_feature_ is passed through a multilayer perceptron (MLP) to produce δ_1_, which encodes the mutation-induced change based solely on local sequence embeddings.

Finally, δ_1_ and δ_2_ are concatenated and fed into a final MLP, which predicts the ΔΔG value of protein point mutation. This hierarchical design enables ProStab to jointly capture both local sequence perturbations and their broader structural consequences, thereby improving the accuracy of stability predictions.

### Benchmark performance of ProStab on various datasets

In this section, we systematically benchmark ProStab on ten independent test sets after training on the Megascale-train split (see *Materials and Methods* for details). These datasets span a wide range of protein families, mutation types, and stability measurement protocols, providing a robust benchmark for assessing distributional generalization. Fig. 2a quantifies the average number of mutants per protein across datasets, revealing a stark contrast between the densely annotated Megascale splits (∼900–1,000 mutants/protein) and conventional DMS datasets such as Fireprot or S669 (as low as 7 mutants/protein), highlighting the challenge of generalizing from sparse mutational coverage. Fig. 2b further contextualizes dataset scale in terms of protein and mutant count. Complementarily, Fig. 2c compares protein length distributions, showing that Megascale predominantly consists of compact proteins (∼70 residues), whereas test sets like S2648 contain many longer sequences, introducing additional generalization difficulty due to increased structural and mutational complexity. We assessed performance using Spearman’s rank correlation coefficient (SCC) to capture ranking consistency and Pearson’s correlation coefficient (PCC) to evaluate linear predictive accuracy.

**Figure 2.**
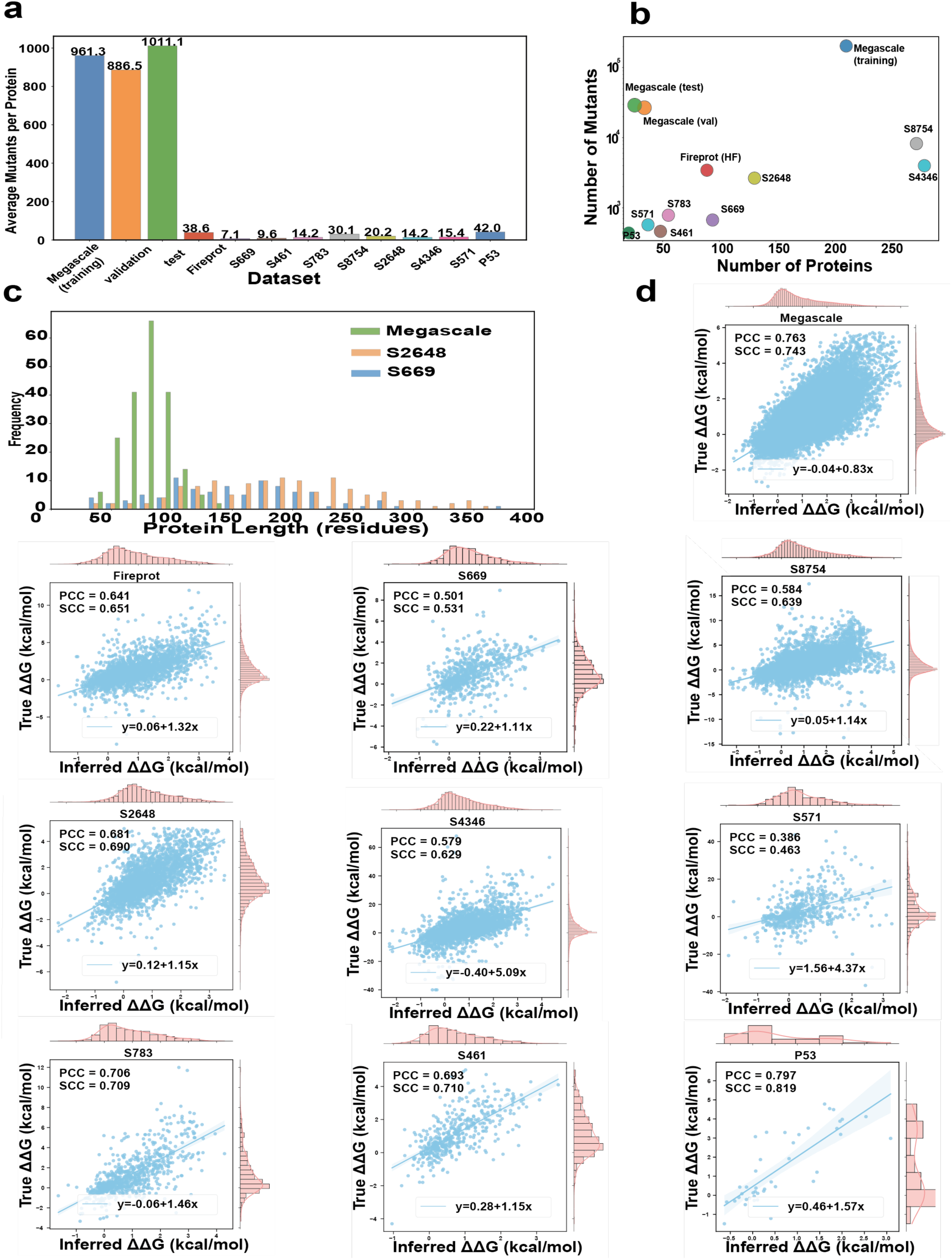
Benchmark performance of ProStab on ten independent test datasets. (a) Average number of mutants per protein across all datasets. (b) Comparison of dataset scale in terms of the number of proteins (x-axis) and total number of mutants (y-axis, log scale). (c) Distribution of protein lengths for three representative datasets. (d) Each scatter plot shows predicted versus experimentally measured ΔΔG values, along with the corresponding regression line and performance metrics: Pearson correlation coefficient (PCC) and Spearman’s rank correlation coefficient (SCC).

Fig. 2d illustrates the distribution of predicted versus experimentally measure ΔΔG values. The test datasets vary markedly in their ΔΔG value distributions, particularly in dynamic range. For example, Megascale spans a narrow ΔΔG range (−3 to 5 kcal/mol), whereas S8754 covers a much broader range (−14 to 15 kcal/mol). This discrepancy limits the generalization ability of models trained on narrow-range datasets like Megascale when applied to broader-range targets. Similarly, for S571 and S4346 — which assess changes in dissolution temperature (ΔΔTm)—the predicted range (−2 to 4) is considerably narrower than the ground truth range (−40 to 60), indicating large absolute deviations. Despite this, ProStab preserves the overall ranking trend, achieving SCC values of 0.463 and 0.629 on S571 and S4346, respectively. These results indicate that ProStab generalizes more effectively than existing methods, particularly across distribution shifts. However, its accuracy in absolute ΔΔG prediction declines when extrapolating beyond the dynamic range of the training data. This limitation likely arises from the narrow span of the Megascale dataset, which may bias the model toward moderate-range outputs and underrepresent extreme stability shifts.

### Performance comparison with state-of-the-art models

We further compared ProStab with a suite of representative baseline methods, including physics-based models (e.g., FoldX^11^, Rosetta^12^) and deep learning-based models (e.g., SPURS^21^, ThermoMPNN^6^, StabilityOracle^18^, Pythia^20^).

As shown in Fig. 3a, ProStab achieved state-of-the-art SCC performance on 7 out of 10 test sets and remained competitive on the remaining three sets. On an extensively used independent test set S669, ProStab outperformed all other methods by substantial margin. For PCC, shown in Fig. 3b, ProStab attained comparable or superior performance across most datasets. These results highlight ProStab’s strong ranking fidelity and stable linear predictive power across diverse mutational and thermodynamic contexts.

**Figure 3.**
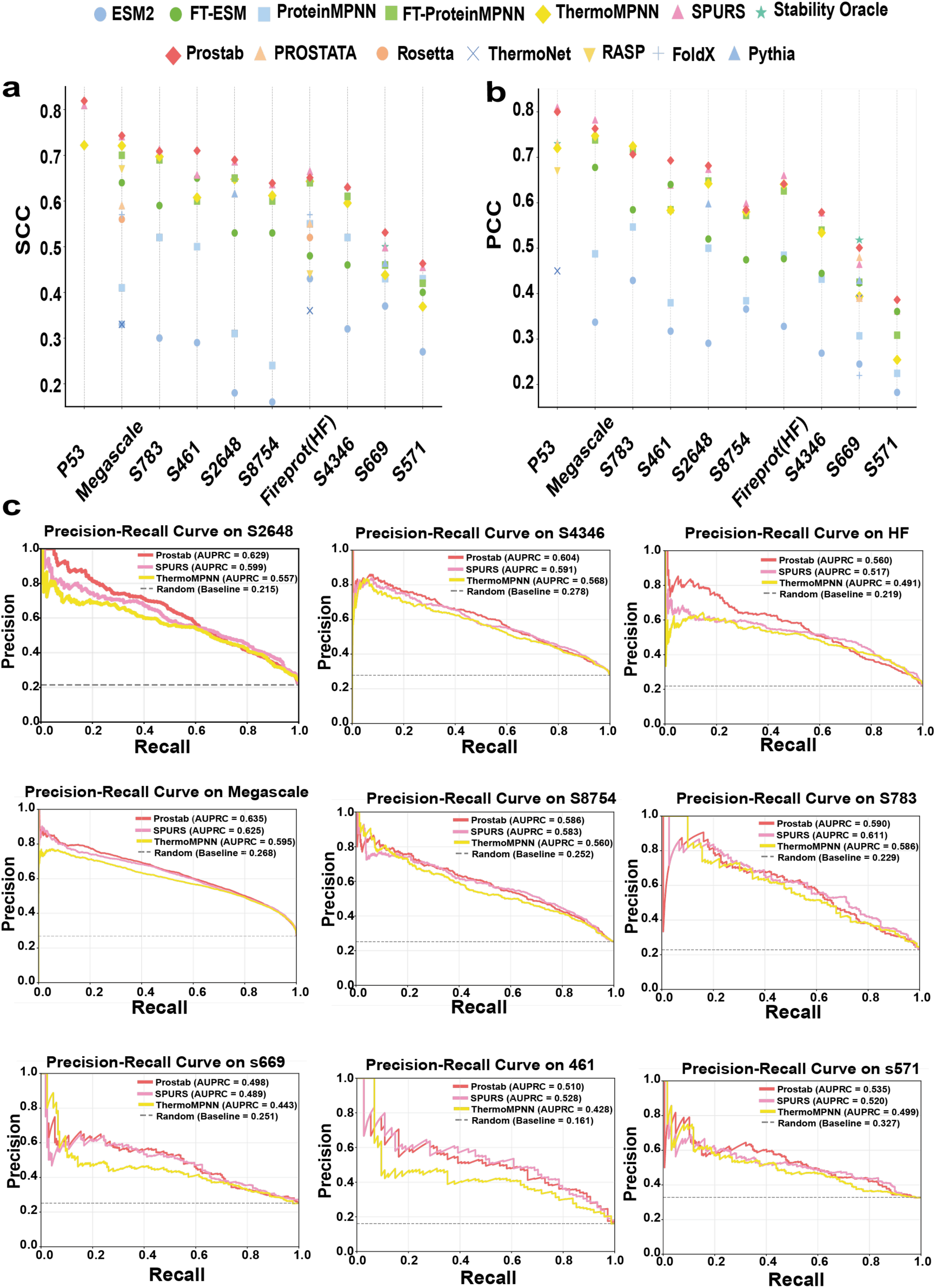
Comparative performance of ProStab and representative baseline models across ten test datasets. (a-b) SCC and PCC comparison, each point represents a model–dataset pair. (c) Precision–recall curves for ProStab and two SOTA methods on 9 out of 10 benchmark datasets (Results for the remaining dataset, P53, are provided in Supplementary Materials).

In addition to PCC and SCC, we further assessed model performance using the area under the precision–recall curve (AUPRC), a metric particularly suited to imbalanced classification settings. In protein stability datasets, stabilizing mutations (positive class) are typically much rarer than destabilizing ones, making AUPRC more informative than AUROC for evaluating a model’s ability to identify rare, functionally desirable variants. For each test set, we computed AUPRC values for ProStab and two strong baselines: ThermoMPNN and SPURS. As shown in Fig. 3c, the dashed horizontal line in each plot indicates the expected AUPRC of a random classifier, which corresponds to the fraction of stabilizing mutations in that dataset—i.e., the baseline level of precision given the positive class prevalence. Across all ten benchmarks, ProStab consistently matches or outperforms both baselines in AUPRC. It achieves the highest scores on datasets such as S2648 (0.629), Megascale (0.635), and S8754 (0.586). These results underscore ProStab’s robust discriminative capability in identifying stabilizing mutations across diverse and skewed mutational landscapes.

### Ablation study and impact of structure quality

We conducted ablation experiments on individual components of ProStab. Fig. 4a and 4c show changes in SCC and PCC observed after removing specific features or components. The full ProStab model consistently outperforms all ablated variants across most datasets. Notably, removing the Transformer Encoder or the fusion feature leads to a significant performance drop, underscoring the crucial role of sequence– structure integration. Unexpectedly, removing δ_feature_ yielded comparable or even improved performance on certain datasets. Its impact appears to be dataset-dependent: its removal produces the lowest performance on the S669 dataset but improves results on Megascale. This discrepancy likely stems from differences in protein length, mutation distribution, or structural complexity. Nonetheless, incorporating δ_feature_ generally improves performance across the majority of datasets.

**Figure 4.**
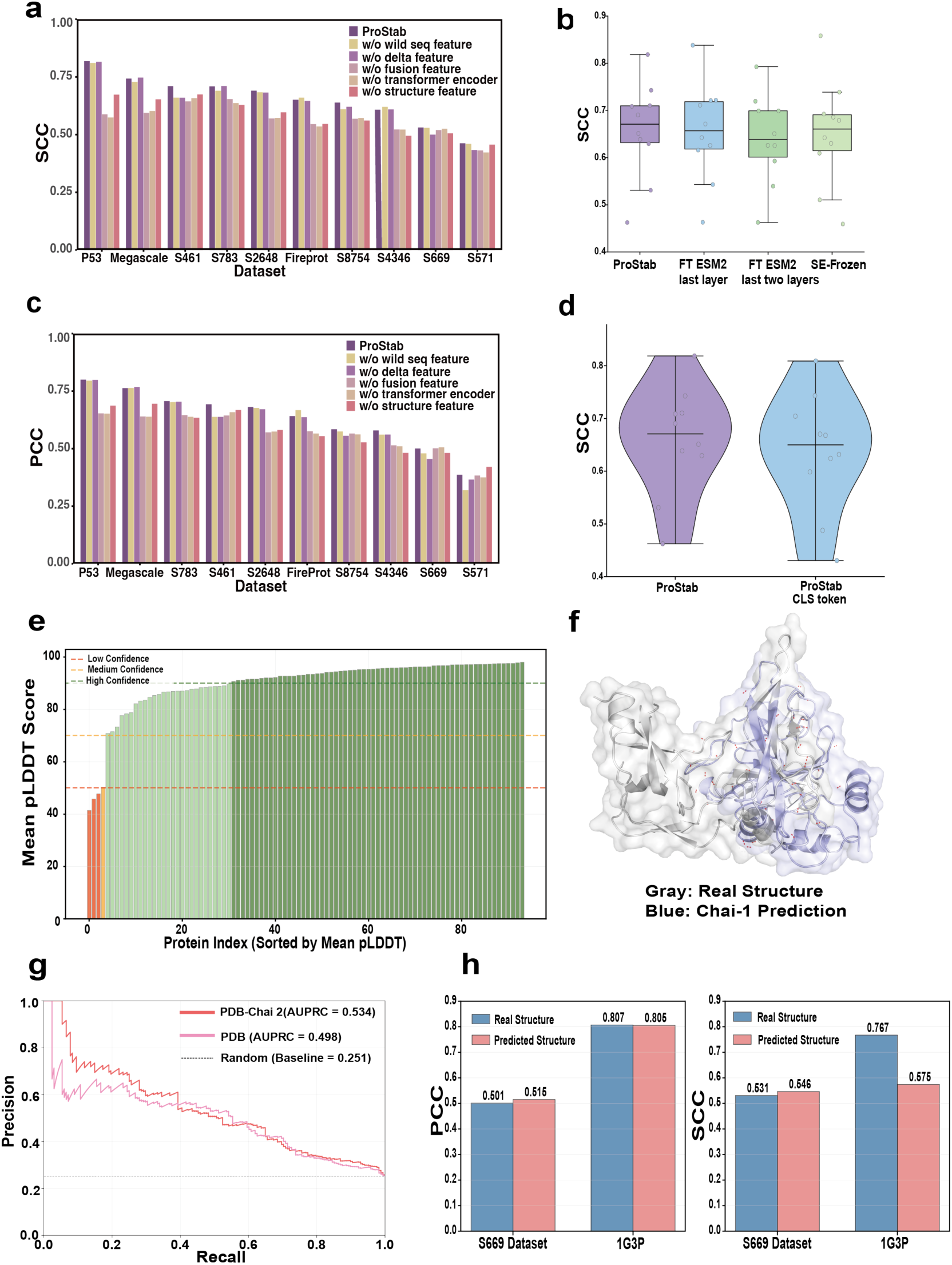
Model ablation and performance under predicted vs. experimental structures. (a) SCC of ProStab and its ablated variants. (b) SCC under different ESM2 fine-tuning strategies, alongside evaluation of parameter freezing in the structure encoder (SE-Frozen). (c) PCC of ProStab and its ablated variants. (d) Comparison between the proposed δ_feature_ representation and the CLS token difference method for modeling mutation effects. (e) Distribution of average predicted pLDDT scores for each of the 94 proteins in the S669 dataset, as predicted by Chai-1. (f) Structural alignment of experimental structure (gray) and Chai-1 prediction (blue) for PDB ID 1G3P; red dashed lines indicate regions exhibiting significant deviations (>2.0 Å). (g) Precision– recall curves computed using experimental structures and predicted structures on the S669 dataset. (h) PCC and SCC computed using experimental and predicted structures for the entire S669 dataset and for the protein 1G3P, which exhibited the lowest average predicted pLDDT score.

We evaluated the impact of different ESM2 fine-tuning strategies on ProStab’s predictive performance. Specifically, we compared the default ProStab configuration—using a frozen ESM2 model—with two variants: fine-tuning the final transformer layer (FT ESM2 last layer) and the last two layers (FT ESM2 last two layers) (Fig. 4b). Among these, the default ProStab achieves the highest median SCC and demonstrates the most stable performance across datasets. Fine-tuning the final layer leads to a slight performance decline, while tuning the last two layers results in further degradation. We also assessed the impact of freezing the structure encoder (SE-Frozen). This configuration yielded the poorest performance overall, highlighting the importance of jointly optimizing the structural encoder for robust ΔΔG prediction.

In addition to model-level tuning, we also examined alternative strategies for encoding mutation-induced differences. Specifically, we replaced the position-specific δ_feature_ with the difference between the [CLS] tokens of the wild-type and mutant sequences. As shown in Fig. 4d, the δ_feature_ yields a slightly higher median SCC and lower variance across datasets. These results indicate that while the [CLS] token captures global sequence context, residue-level features provide a more stable and discriminative signal for modeling mutation effects.

To investigate the model’s sensitivity to predicted versus experimental structures, we employed Chai-1^33^ to predict structures for all 94 proteins involved in the S669 dataset. The average pLDDT score across these proteins was 89.9, indicating high overall prediction quality (Figure 4e). As a case study, we selected protein 1G3P, which exhibited the lowest average pLDDT score (41.5) and contained 27 mutation sites in the S669 dataset. Figure 4f shows the structural alignment between the experimental structure (gray) and Chai-1 prediction (blue) for 1G3P, with red dashed lines indicating regions of significant deviation (>2.0 Å). Surprisingly, the model demonstrated superior performance on predicted structures compared to experimental structures. As illustrated in Figure 4g, the AUPRC achieved using predicted structures on the S669 dataset reached 0.534, while that using experimental structures was only 0.498. Figure 4h further presents detailed performance comparisons on both the S669 dataset and the 1G3P protein. On the S669 dataset, predicted structures showed marginally better performance than experimental structures in both PCC and SCC metrics, though the differences were relatively modest. For the 1G3P case study, both structures exhibited comparable PCC performance, while the predicted structure yielded a lower SCC (0.575) compared to the experimental structure (0.767). This counterintuitive result can be attributed to several factors. First, the training dataset Megascale contains a proportion of predicted structures, enabling the model to learn and adapt to the systematic features and bias patterns inherent in predicted structures during training. Second, predicted structures may exhibit a “denoising” effect; compared to experimental structures that may contain local distortions due to crystal packing effects, solvent conditions, and other experimental artifacts, predicted structures generated based on evolutionary constraints and physical principles may better represent the functional conformations of proteins. Finally, our model likely relies primarily on large-scale structural features such as secondary structure and domain organization rather than precise atomic coordinates for prediction, making the high consistency between predicted and experimental structures at these coarse-grained feature levels sufficient to support robust predictive performance.

### Evaluating ProStab’s ability to identify stabilizing mutations

To evaluate ProStab’s ability to identify stabilizing mutations, we compared its performance with SPURS^21^ and ThermoMPNN^6^ on the S2648, S783, and S669 datasets under various ΔΔG thresholds. Fig. 5a summarizes precision and recall across these datasets. On S2648, ProStab consistently maintains high precision across ΔΔG thresholds, reaching 0.9 at the −1.0 kcal/mol threshold. On S669, it outperforms both baselines at stricter thresholds while performing comparably at milder ones. Meanwhile, ThermoMPNN achieves the highest precision on S783. In terms of recall, SPURS performs best across most datasets and thresholds. These results suggest that ProStab provides high-confidence predictions with relatively limited coverage, making it a practical choice in scenarios with constrained experimental validation capacity.

**Figure 5.**
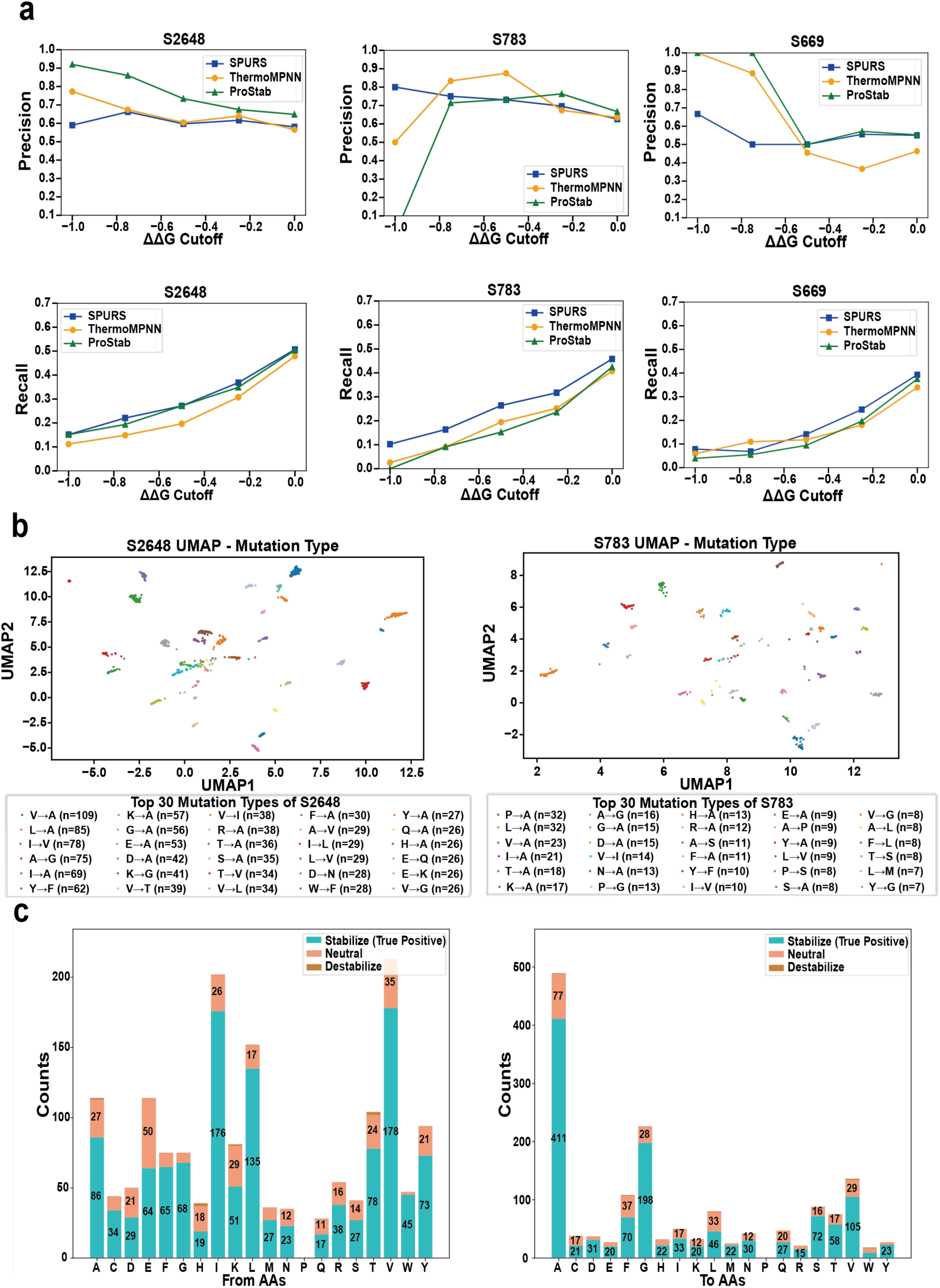
Benchmarking performance, feature representation, and amino acid substitution patterns across datasets. (a) Precision and recall for ProStab, SPURS, and ThermoMPNN on the S2648, S783, and S669 datasets under varying ΔΔG threshold cutoffs. (b) UMAP projections of the δ_feature_ for mutations in the S2648 and S783 datasets, illustrating the learned mutation-level representations. (c) Distribution of experimentally observed stabilizing, neutral, and destabilizing mutations in the S2648 dataset, grouped by source (from) and target (to) amino acid types. All mutations shown are from the experimentally validated set of stabilizing substitutions. Mutations correctly predicted by ProStab as stabilizing (ΔΔG < –0.5 kcal/mol) are highlighted in green.

To investigate the structure of mutation-induced embeddings, we visualized δ_feature_ using UMAP (Uniform Manifold Approximation and Projection for Dimension Reduction) on the S2648 and S783 datasets (Fig. 5b). Each point represents a mutation, with colors denoting specific amino acid substitution types. The projection shows that mutations of the same substitution type tend to cluster together in δ_feature_ space, indicating that the pretrained embeddings encode substitution-specific patterns in a consistent and structured manner.

We further assessed ProStab’s prediction consistency across amino acid contexts by analyzing its performance on the S2648 dataset, grouped by source (From AAs) and target (To AAs) residues (Fig. 5c). Stabilizing mutations were defined based on experimental ΔΔG values (ΔΔG < −0.5 kcal/mol), and only these were included in the analysis. Each bar shows the total number of experimentally stabilizing mutations for a given amino acid, while the green segment represents those also predicted by the model as stabilizing (i.e., predicted ΔΔG < −0.5 kcal/mol). Predictions were additionally categorized as neutral (|ΔΔG| ≤ 0.5 kcal/mol) or destabilizing (ΔΔG > 0.5 kcal/mol)^18^. These results indicate that ProStab correctly identifies a substantial proportion of stabilizing mutations across a wide range of amino acid substitutions. Notably, this trend holds across both source and target residues, highlighting ProStab’s robustness in detecting stabilizing effects across diverse mutation directions.

### Evaluation Across Physicochemical Mutation Categories

To assess how ProStab and SPURS perform across mutations involving different physicochemical property transitions, we categorized amino acids based on their side-chain characteristics—specifically, hydrophobic, hydrophilic, and neutral residues, as well as positively charged, negatively charged, and uncharged groups. We then evaluated the change in prediction error (ΔRMSE) between the two models by mutation type (Fig. 6a). A negative ΔRMSE indicates better performance of ProStab, while a positive value favors SPURS. Results on additional benchmark datasets are provided in Supplementary Figures.

**Figure 6.**
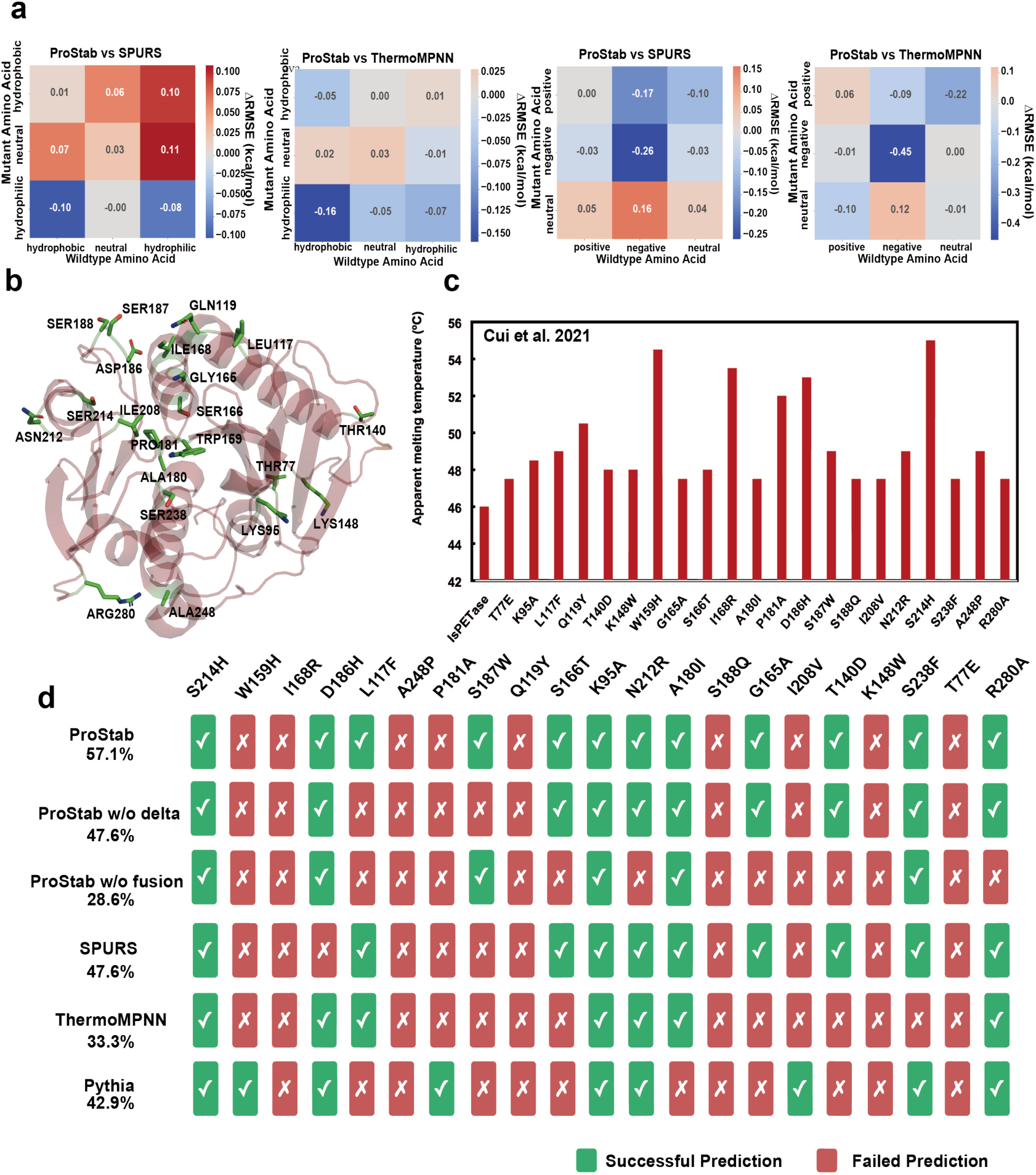
Error analysis on the S783 dataset and experimental validation on IsPETase. (a) Differential prediction error (ΔRMSE, kcal mol⁻¹) between ProStab and baseline models (SPURS, ThermoMPNN), grouped by wildtype and mutant amino acid polarity or charge. Negative values indicate improved accuracy by ProStab. (b) Structural mapping of 21 validated single-point mutation sites on IsPETase. (c) Apparent melting temperatures (°C) of single-point mutations in IsPETase^34^. (d) Prediction accuracy of ProStab and its ablated variants compared to baseline methods (SPURS, ThermoMPNN, and Pythia) on 21 known stabilizing mutations in IsPETase; green represents successful predictions and red represents failed predictions.

When benchmarked against ThermoMPNN, ProStab consistently outperformed it across most evaluated conditions. The most pronounced improvements were observed for mutations resulting in hydrophilic amino acids. Further analysis based on wildtype charge states revealed that ProStab conferred clear advantages across most charge categories, with particularly strong improvements for negatively charged wildtype residues. The largest individual gain was observed for negative wildtype to negative mutant substitutions, achieving a ΔRMSE of −0.45. In comparison to SPURS, ProStab offered only modest gains in select hydrophobicity-defined categories, with |ΔRMSE| typically ≤ 0.10 kcal mol⁻¹. Consistent with the results against ThermoMPNN, a significant improvement was likewise observed in the comparison with SPURS for negative wildtype to negative mutant substitutions, with a ΔRMSE of −0.26.

### Evaluation on Beneficial Mutations in IsPETase Engineering

We conducted a case study on the thermostability engineering of IsPETase to assess ProStab’s ability to identify rare stabilizing mutations. Specifically, we used 21 single-point mutations previously reported to increase its melting temperature (Fig. 6c) as an external benchmark^34^. Fi. 6b shows the structural mapping of these mutation sites on the IsPETase structure.

ProStab achieved the highest success rate, correctly identifying 12 of 21 beneficial mutations (57.1%), outperforming SPURS (10/21, 47.6%), Pythia (9/21, 42.9%), and ThermoMPNN (7/21, 33.3%) (Fig. 6d). To further evaluate the contributions of δ_feature_ and the fusion feature, we performed ablation experiments on this benchmark. Removing δ_feature_ reduced the number of correct predictions to 10, while excluding the fusion feature led to a more substantial drop to 6. These results highlight the complementary roles of both components: the δ_feature_ captures localized sequence perturbations, while the fusion feature integrates sequence and structural context—both essential for accurately identifying stabilizing mutations.

### Attention Weights Reveal Model Sensitivity to Protein Structural Interactions

To further investigate the interpretability of ProStab, we analyzed how its attention mechanism captures key structural interactions within proteins. Specifically, we visualized attention weights from the Transformer encoder to examine whether the model prioritizes reside-residue contacts that align with known structural interactions. In Fig. 7a, we examined residue TYR17 in protein 6OBK. The attention heat map revealed strong associations between TYR17 and positions 5, 6, 12, and 13. Structural visualization confirmed them as non-covalent contacts (distance < 3.5 Å) with ILE5, LEU6, LEU12 and THR13. To assess the model’s sensitivity to structural changes, we used FoldX to generate a mutant structure in which TYR17 is replaced by glycine. The resulting attention distribution changed markedly, with reduced focus on the original contact residues—corresponding to weakened interactions in the mutated structure. A similar analysis was conducted for TYR3 in the GB1 domain (PDB: 3GB1; Fig. 7b). In the wild-type structure, TYR3 forms notable contacts with ASP22, ALA26, and LEU5. Upon mutation to alanine, attention was redistributed, likely reflecting the loss of the aromatic side chain. In Fig. 7c, we further analyzed attention patterns in three additional proteins—2KWH, 1A32, and 1B71—focusing on residue HIS21, ARG40, and TYR63, respectively. In each case, the model assigned high attention scores to known interaction partners: LEU17, LEU20, and LEU24 (2KWH), ILE5, ILE6, AND ILE13 (1A32); and VAL30 and ILE32 (1B71).

**Figure 7.**
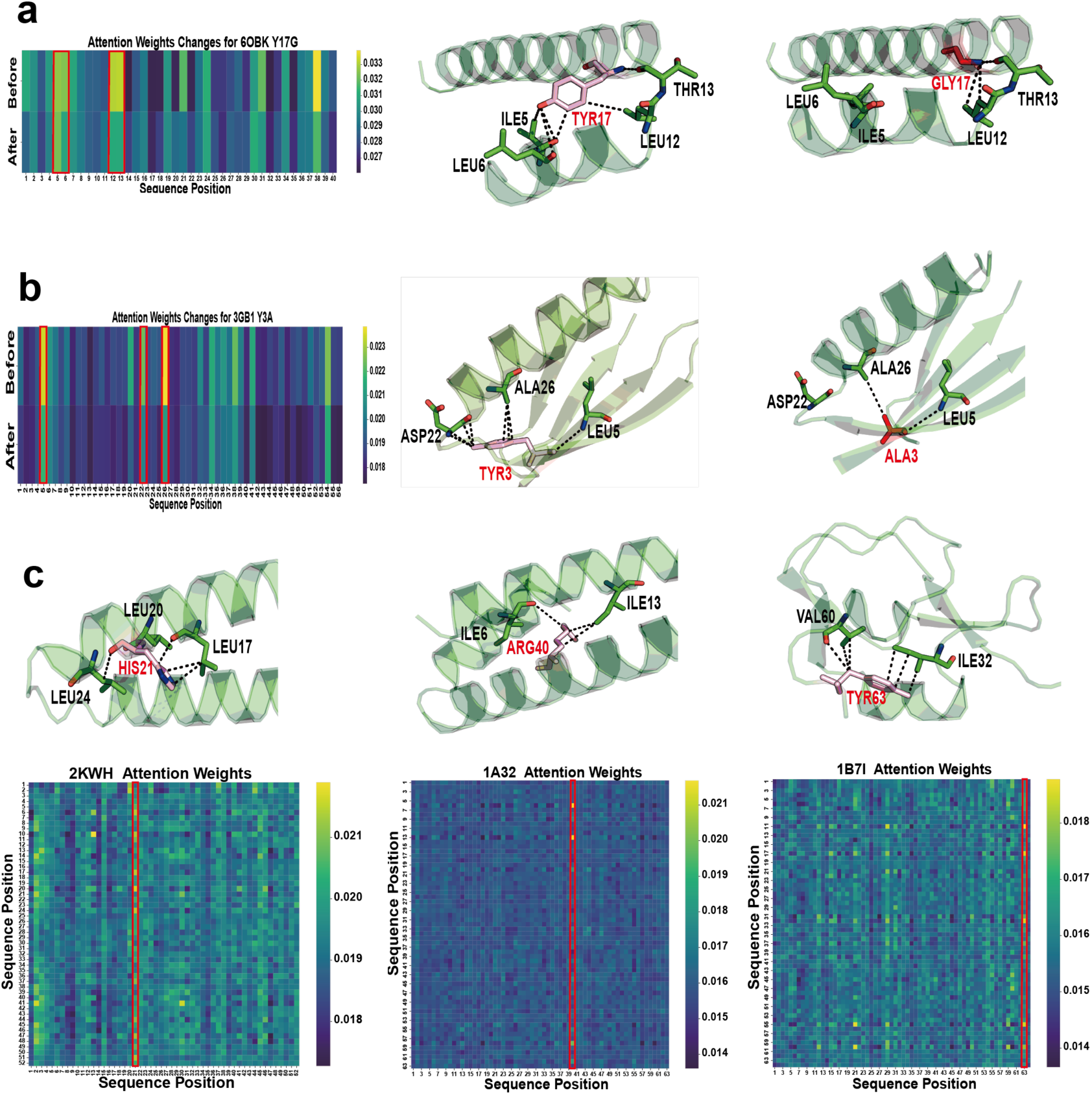
Attention mechanisms capture residue-level structural interactions. (a) Attention analysis of the TYR17→GLY mutation in the 6OBK protein. Left: heat map showing changes in attention weights before and after the mutation. Right: structural visualization illustrating the interaction between TYR17 (red) and neighboring residues ILE5, LEU6, LEU12, and THR13 (green), as well as the disruption of these interactions following the substitution with GLY. (b) Attention analysis of the TYR3 → ALA mutation in the 3GB1 protein. Left: heat map of attention weights before and after the mutation. Right: structural visualization of TYR3 and its interactions with ASP22, ALA26, and LEU5, highlighting changes upon mutation. (c) Attention analysis of representative key residues across multiple proteins. Top row: structural interactions of HIS21 in 2KWH, ARG40 in 1A32, and TYR63 in 1B71 (all shown in red) with their local environments. Bottom row: corresponding attention heat maps, where brighter regions indicate higher attention values. In all structures, dotted lines represent non-covalent interactions within 3.5 Å.

These findings suggest that ProStab effectively captures biologically meaningful intramolecular interactions and dynamically responds to structural perturbations. The ability to reflect physical residue–residue contacts—based solely on learned attention over input features—enhances the interpretability and reliability of ProStab’s predictions, distinguishing it from models that rely primarily on sequence-level correlations.

### Identification of stabilizing mutations for an ATA

Encouraged by the excellent predictive performance of ProStab on ΔΔG estimations, we experimentally validated its predictions using the amine transaminase from Exophiala xenobiotica (Fig. 8a). Based on the model outputs, top 20 mutations were introduced across 10 surface-exposed residues (Fig. 8b), all distal to the enzyme’s active site—consistent with the empirical observation that stabilizing mutations often occur at peripheral regions. Among these, 16 variants were solubly expressed in Escherichia coli, and four demonstrated markedly improved thermostability compared to the wild-type enzyme (Fig. 8c). Notably, two of the top five predicted variants exhibited substantial enhancements, with the best-performing mutant (A245I) retaining 99.6% of catalytic activity after thermal challenge, compared to just 27.3% for the wild-type (Fig. 8d). Fig. 8e and 8f demonstrate that the mutations significantly enhance the enzyme’s stability at elevated temperatures and during prolonged heat treatment at 40°C. These results underscore the practical utility of ProStab in efficiently prioritizing stabilizing mutations and reducing experimental screening efforts.

**Figure 8.**
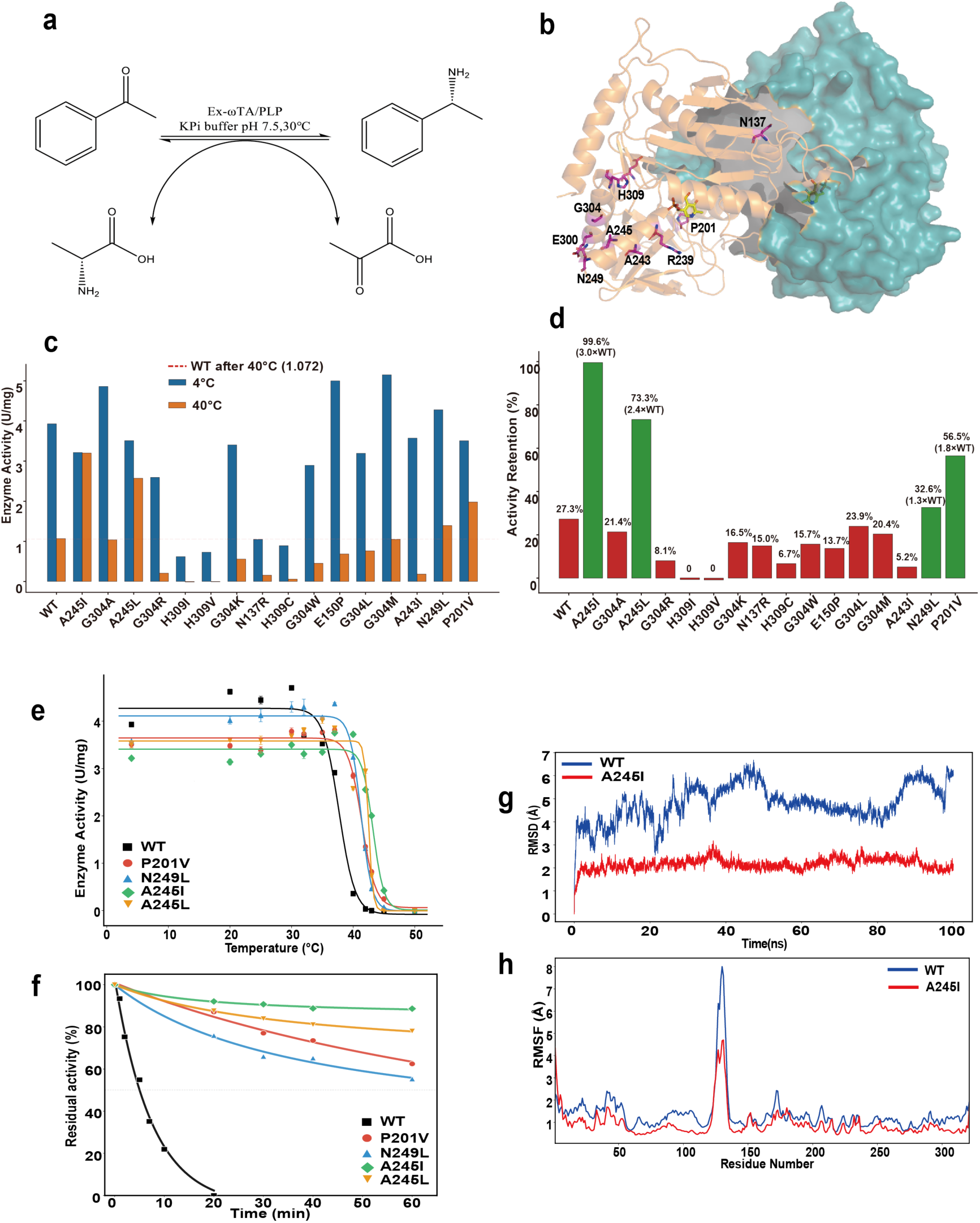
ProStab-predicted mutations in Ex-ATA enhance thermostability, as validated by wet-lab experiments and molecular dynamics simulations. (a) Schematic representation of the transamination reaction catalyzed by Ex-ATA in the presence of PLP (pyridoxal 5’-phosphate) in phosphate buffer (pH 7.5, 30°C). (b) Crystal structure of Ex-ATA highlighting the 10 residues selected for mutation based on ProStab predictions (shown in magenta sticks). The enzyme dimer is shown with one monomer in cartoon and the other in surface representation. (c) Enzyme activity of wild-type (WT) and variant Ex-ATA measured before and after heat treatment at 40°C for 1 hour. Blue bars indicate activities measured at 4°C, and orange bars show activities post-incubation at 40°C. The red dashed line represents the residual activity level of WT after heat treatment. (d) Residual activity retention (%) after heat treatment at 40°C. Activity retention is normalized to untreated WT levels. Four variants (A245I, A245L, N249L, and P201V) exhibited significantly improved thermostability compared to WT. (e) Thermal deactivation of selected variants and WT at various temperatures for 10 minutes. (f) The thermal deactivation half-life of WT and the variants at 40°C. (g) Molecular dynamics simulation of WT and A245I mutant showing backbone RMSD over a 100 ns trajectory. A245I showed reduced global fluctuation, indicating increased structural rigidity. (h) Per-residue RMSF analysis of WT and A245I mutant. The mutation site region (around residue 245) and a distal flexible region (around residue 130) both exhibited dampened dynamics in the A245I variant, suggesting potential long-range stabilization effects.

To elucidate the molecular basis of the observed stability enhancement, we performed molecular dynamics (MD) simulations comparing the wild-type enzyme and the A245I variant. The mutant displayed a significantly reduced root-mean-square deviation (RMSD) over the simulation trajectory (Fig. 8g), indicative of a globally more rigid structure. Residue-wise root-mean-square fluctuation (RMSF) analysis further revealed decreased flexibility around the mutation site (Fig. 8h), supporting a stabilization mechanism via entropy reduction. Interestingly, the A245I variant also exhibited suppressed dynamics at a distal region near residue 130—an intrinsically flexible segment—suggesting a potential long-range dynamic coupling. Such effects may propagate through structural or energetic networks, highlighting how single-point mutations can modulate nonlocal conformational behavior.

### Deployment of an Online Interface to facilitate protein stability prediction

To enhance the accessibility and usability of ProStab, we developed a web-based interface for predicting protein stability changes (ΔΔG) induced by single-point mutations, available at https://ails.sjtu.edu.cn/prostab. Users can upload a protein structure (PDB format), specify the chain ID, and define mutations by entering wild-type residue, its position, and the substituted amino acid. Upon submission, the model computes the ΔΔG using the full ProStab framework and returns both the prediction ΔΔG value and an intuitive stability warning. For example, in the case of a V1A mutation in PDB entry 1A0N (chain A), the predicted ΔΔG is 0.12 kcal/mol, and the substitution is flagged as potentially destabilizing. This interactive platform enables practical ΔΔG estimation without requiring local setup or coding expertise, offering an accessible entry point for researchers in protein engineering, stability optimization, and mutational scanning.

## Discussion and Conclusion

In this study, we introduced ProStab, a deep learning framework for predicting the change in protein thermal stability (ΔΔG) upon single-point mutations. Distinct from existing methods^21,35–37^, ProStab integrates mutation-specific sequence perturbations and wild-type structural priors without relying on explicit mutant structure prediction. Across multiple benchmark datasets, ProStab consistently achieves state-of-the-art performance in ΔΔG regression and classification tasks.

Beyond overall performance, ProStab demonstrates strong predictive power for stabilizing mutations. Under varying ΔΔG thresholds, it achieves higher accuracy than baseline models, and in the case study on IsPETase, it identifies the largest number of beneficial mutations among all competing methods. To assess real-world applicability, we conducted experimental validation on the transaminase, where 4 out of 16 tested mutations predicted by ProStab improved thermal stability. Notably, the 1st top-ranked prediction yielded the highest experimental stability, highlighting ProStab’s practical utility in protein engineering. A publicly available web server has also been developed to support high-throughput use.

Despite these promising results, ProStab currently does not utilize structural models of the mutant protein. Although recent tools such as AlphaFold2^38^ have advanced single-sequence structure prediction, their mutation-level sensitivity remains limited. This is largely due to the reliance on multiple sequence alignments (MSAs) and template reuse, which often produce mutant structures nearly identical to the wild type for point mutations. Furthermore, some single-point mutations may not significantly alter static structures but instead affect dynamic properties, such as flexibility or conformational transitions, which are difficult to model due to limited experimental data and the high computational cost of molecular dynamics simulations.

Another limitation lies in the handling of higher-order (multi-point) mutations, which ProStab does not yet address. Extending to this setting poses two main challenges: (1) the lack of large-scale training data covering combinatorial mutation space, and (2) the need for models to capture complex epistatic interactions and cumulative effects across multiple residue substitutions^39–41^. These directions represent an important focus for future work.

In summary, ProStab offers a scalable, accurate, and structure-aware framework for ΔΔG prediction of single-point mutations. While challenges remain in dynamic modeling and generalizing to multi-mutant contexts, ProStab represents a meaningful step toward practical and intelligent protein design.

## Materials and methods

### Datasets

The Megascale dataset is a high-throughput protein folding stability dataset generated via cDNA display proteolysis. In this study, we partitioned the Megascale dataset into training, validation and test sets following the same protocol used in SPURS^21^. The training and validation sets were used for model training and hyperparameter optimization. In addition to the Megascale test set, we employed model performance on nine independent benchmark datasets. All evaluation sets exhibit low sequence similarity (<25%) to the training set, ensuring a robust assessment of generalization. Detailed statistics for each dataset are provided in the Supplementary Information.

### Training settings

ProStab was trained using the AdamW optimizer (learning rate = 0.0001, betas = [0.9,0.98], weight decay = 0.01). The model was optimized with the mean squared error (MSE) loss for up to 50 epochs. Each training batch contained a single protein (batch size = 1), and all experiments were conducted on a single NVIDIA RTX 4090 GPU. Model selection was based on the highest SCC achieved on the validation set, with early stopping applied (patience = 20). A plateau learning rate scheduler was used to reduce the learning rate by a factor of 0.2 if the validation RMSE plateaued.

### PLM-Based Representation and Mutation Feature Construction

We extract residue-level protein representations using the pretrained ESM2-650M protein language model. Given a wild-type protein sequence of length *L*, the model outputs an embedding of shape *L* × 1280 after removing the special [CLS] and end-of-sequence tokens. we then obtain the full-sequence embeddings for both the wild - type and mutant sequences:

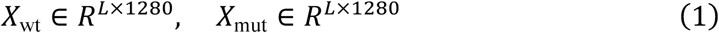

For a mutation occurring at position *i*, we capture its localized effect by defining a delta feature as the difference between the embeddings of the wild-type and mutant residues at that position:

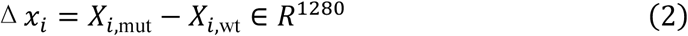

In parallel, the full-sequence embedding of the wild-type protein, *X*_wt_, is fused with the corresponding structural features *S* ∈ *R^L^*^×*d*)^ extracted from a structure encoder:

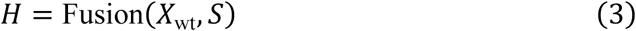

where Fusion(⋅) denotes the feature integration module, as described in Section 3.4.

### Graph-Based Structure representation

ProStab integrates ProteinMPNN as its structural encoder. ProteinMPNN is a graph-based message passing neural network (MPNN) designed to learn residue-level representations from the three-dimensional structures of proteins. It models each protein as a graph *G* = (*V*, *E*), where *V* represents nodes corresponding to backbone atoms (N, Cα, C, O, and Cβ), and E denotes edges connecting each atom to its 48 nearest neighbors. This graph construction enables the encoder to effectively capture local structural topology and geometric context.

The final output of the structural encoder is denoted as:

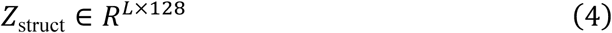

By default, ProteinMPNN adopts a random autoregressive decoding strategy, in which only masked positions are predicted. The decoding order is determined by assigning random values to each residue and sorting them, resulting in a unique decoding sequence for each sample. Under this framework, the prediction for residue *i* is conditioned on its structural embedding and the amino acids decoded at prior positions, and can be modeled as:

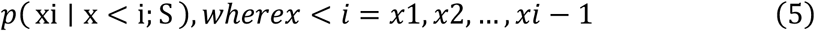

While this strategy is effective for generative tasks, it limits the model’s ability to capture global sequence context. To overcome this limitation, ProStab adopts a non-autoregressive one-shot decoding strategy, which enables the model to better leverage the full structural environment.

Under this setting, the prediction at position *i* is conditioned on the protein structure and all other amino acids in the sequence except for the target residue. This can be formulated as:

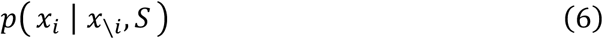

Where *x*_∖*i*_ denotes all residues in the sequence except *x_i_*.

To implement the one-shot decoding mechanism, we introduce a visibility mask matrix *M* ∈ {0,1}*^L^*^×*L*^, defined as:

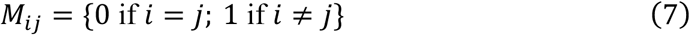

This mask ensures that all residues except the target position are visible during prediction, allowing the model to utilize full-sequence context and better capture structural features relevant to protein stability.

To support this decoding scheme, we fine-tune ProteinMPNN in an end-to-end manner, enabling the structural encoder to adapt its representations to the stability prediction task.

In addition, we extract and retain the intermediate decoder outputs from each of the N=3 decoder layers for all positions:

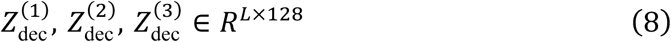

We also incorporate a residue identity embedding, *Z*_id_ ∈ *R^L^*^×128^, obtained by mapping each amino acid type to a 128-dimensional vector via the embedding layer of ProteinMPNN.

Finally, the residue identity embedding is concatenated with the decoder outputs to form the final structure representation:

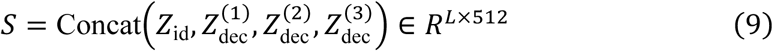

### Sequence–Structure Feature Fusion via a Transformer Encoder

To enable joint modeling of sequence and structural contexts, we fuse the full-sequence representation of the wild-type protein with residue-level structural features. These two modalities are concatenated along the feature dimension to form the initial fused representation:

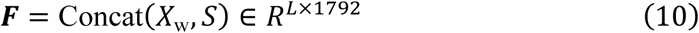

This fused matrix *F* is then processed by a two-layer Transformer encoder, which enables interaction and contextualization between sequence and structure across all residues.

Formally, the final integrated representation is computed as:

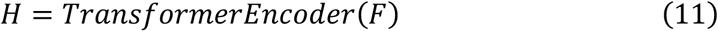

The Transformer encoder is configured with a hidden size of 1792, 8 attention heads, a feed-forward network dimension of 3584, and a dropout rate of 0.1.

### ΔΔG Prediction via Dual-Component Modeling

To accurately model protein stability changes upon mutations, we adopt a dual-component prediction strategy that explicitly separates two sources of information:

1. Mutation-induced sequence perturbation—representing the actual difference introduced by the mutation.
2. Structure-informed susceptibility to mutation—derived entirely from the wild-type sequence and structure.

### Mutation-Induced Delta Feature (δ₁)

As previously defined, the mutation-induced delta feature at residue position I is computed as:

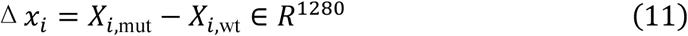

This difference vector is passed through a multi-layer perceptron ℎ: *R*^1280^ → *R* to produce a scalar output:

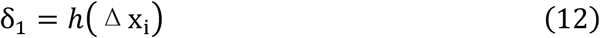

This component captures the direct, mutation-specific sequence-level effect.

### Structure-Informed Stability Projection (δ₂)

In parallel, we utilize the fused sequence–structure representation *H* ∈ *R^L^*^×1792^ from Section 3.4. The embedding at position *i*, *H_i_* is passed through an MLP *g*: *R*^1792^ → *R*^20^, which predicts a vector of ΔG values for all 20 canonical amino acids:

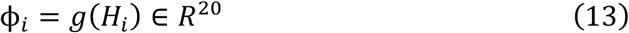

Each element *ϕ_i_*(*a*) represents the predicted stability if residue *i* were mutated to amino acid *a*. Thus, the ΔΔG estimate is then computed as:

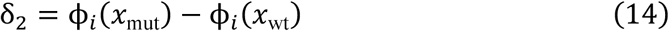

### Final Prediction

The final ΔΔG prediction is obtained by concatenating the two components and passing them through an MLP *f*: *R*^𝟚^ → *R*:

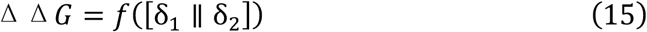

This dual-branch design allows ProStab to simultaneously model mutation-specific perturbations and inferred structural susceptibility, achieving both high accuracy and scalability in protein stability prediction.

### Gene synthesis and site-directed mutagenesis

The E. xenobiotica ATA gene was ordered codon-optimized for expression in E. coli from Hongxun Biotech (Xiamen, China). Mutagenesis was carried out in inverse-PCR using Super-Fidelity DNA polymerase according to the manufacturer’s protocol. Primer list is available in Supplementary Table 1. All plasmids were verified by sequencing.

### Protein Expression and Purification

The E. xenobiotica ATA gene was ordered codon-optimized for expression in E. coli from Hongxun Biotech (Xiamen, China). Plasmids harboring target genes were transformed into E. coli BL21 (DE3) cells for protein production. Cells were grown in LB medium supplemented with 50 μg/ml kanamycin at 37 °C with shaking at 220 rpm until an OD600 of 0.6–0.8 was reached. Protein expression was induced by the addition of 0.1 mM isopropyl-β-d-1-thiogalactopyranoside (IPTG) and the cells were further grown for 20 h at 20 °C with shaking at 220 rpm. After harvesting by centrifugation at 4000 g for 10 min, cell pellets were collected and resuspended and lysed in Lysis Buffer (50 mM potassium phosphate buffer (KPi buffer), pH 7.5, 300 mM NaCl, 20 mM imidazole, 10% TieChui™ E.coli Lysis Buffer (ACE Biotechnology)) at 4℃ for 30min, and centrifuged at 17000 g (60 min, 4 °C) to remove the cell debris. The target protein in the supernatant was purified by nickel-affinity chromatography using gravity-flow columns packed with Ni-NTA resin. The eluted protein was desalted using a desalting column, concentrated by ultrafiltration. Finally, the purified proteins were flash-frozen in liquid nitrogen and stored at −80 °C until use.

### Spectrophotometric Activity Assay

The activity of ExTA and its mutants was monitored by tracking the rise in absorbance at 280 nm for 10 min using a BioTek Synergy H1 microplate reader. The purified wild-type enzyme and mutant enzymes were respectively incubated in a water bath at 20−45 °C for 10 min, and then rapidly placed on ice to cool for 10 min after incubation. The assays were performed at 30 °C in 200 μl reaction mixtures consisting 50 mM KPi buffer (pH 7.5), 0.1 mM PLP, 5 mM (R)-Phenylethylamine, 10 mM Sodium pyruvate and 0.1 mg ml−1 ExTA. The specific enzymatic activity was determined from the rate of absorbance change at 280 nm (ΔA280/min). Each assay was performed in triplicate.

## Acknowledgements

This work was supported by the National Key Research and Development Program of China (Grant Nos. 2024YFA1306902 and 2023YFC2506402), the National Natural Science Foundation of China (Grant No. 62172274), and the Shanghai Municipal Education Commission (Grant No. 2024AIZD008).

## Data Availability Statement

The codes and datasets are open-source and available at https://github.com/xtanh/ProStab.

## References

1. Rahban, M. et al. Thermal stability enhancement: Fundamental concepts of protein engineering strategies to manipulate the flexible structure. Int. J. Biol. Macromol. 214, 642–654 (2022).

2. Goldenzweig, A. & Fleishman, S. J. Principles of Protein Stability and Their Application in Computational Design. Annu. Rev. Biochem. 87, 105–129 (2018).

3. Wu, S., Snajdrova, R., Moore, J. C., Baldenius, K. & Bornscheuer, U. T. Biocatalysis: Enzymatic Synthesis for Industrial Applications. Angew. Chem. Int. Ed. 60, 88–119 (2021).

4. Bell, E. L. et al. Biocatalysis. Nat. Rev. Methods Primer 1, 1–21 (2021).

5. Narayanan, H. et al. Machine Learning for Biologics: Opportunities for Protein Engineering, Developability, and Formulation. Trends Pharmacol. Sci. 42, 151–165 (2021).

6. Dieckhaus, H., Brocidiacono, M., Randolph, N. Z. & Kuhlman, B. Transfer learning to leverage larger datasets for improved prediction of protein stability changes. Proc. Natl. Acad. Sci. 121, e2314853121 (2024).

7. Wang, Y. et al. Directed Evolution: Methodologies and Applications. Chem. Rev. 121, 12384–12444 (2021).

8. Fowler, D. M. & Fields, S. Deep mutational scanning: a new style of protein science. Nat. Methods 11, 801–807 (2014).

9. Arnold, F. H. Design by Directed Evolution. Acc. Chem. Res. 31, 125–131 (1998).

10. Giver, L., Gershenson, A., Freskgard, P.-O. & Arnold, F. H. Directed evolution of a thermostable esterase. Proc. Natl. Acad. Sci. 95, 12809–12813 (1998).

11. Schymkowitz, J. et al. The FoldX web server: an online force field. Nucleic Acids Res. 33, W382–W388 (2005).

12. Kellogg, E. H., Leaver-Fay, A. & Baker, D. Role of conformational sampling in computing mutation-induced changes in protein structure and stability. Proteins Struct. Funct. Bioinforma. 79, 830–838 (2011).

13. Umerenkov, D. et al. PROSTATA: a framework for protein stability assessment using transformers. Bioinformatics 39, btad671 (2023).

14. Savojardo, C., Manfredi, M., Martelli, P. L. & Casadio, R. DDGemb: predicting protein stability change upon single- and multi-point variations with embeddings and deep learning. Bioinformatics 41, btaf019 (2025).

15. Ouyang-Zhang, J., Diaz, D. J., Klivans, A. R. & Krähenbühl, P. Predicting a Protein’s Stability under a Million Mutations. Preprint at 10.48550/arXiv.2310.12979 (2023).

16. Nie, Z. et al. Protein stability prediction with dual-view ensemble learning from single sequence. 2024.04.22.590665 Preprint at 10.1101/2024.04.22.590665 (2025).

17. Li, B., Yang, Y. T., Capra, J. A. & Gerstein, M. B. Predicting changes in protein thermodynamic stability upon point mutation with deep 3D convolutional neural networks. PLOS Comput. Biol. 16, e1008291 (2020).

18. Diaz, D. J. et al. Stability Oracle: a structure-based graph-transformer framework for identifying stabilizing mutations. Nat. Commun. 15, 6170 (2024).

19. Blaabjerg, L. M. et al. Rapid protein stability prediction using deep learning representations. eLife 12, e82593 (2023).

20. Sun, J., Zhu, T., Cui, Y. & Wu, B. Structure-based self-supervised learning enables ultrafast protein stability prediction upon mutation. The Innovation 6, (2025).

21. Rewiring protein sequence and structure generative models to enhance protein stability prediction - PMC. https://pmc.ncbi.nlm.nih.gov/articles/PMC11870403/.

22. Wang, S., Tang, H., Zhao, Y. & Zuo, L. BayeStab: Predicting effects of mutations on protein stability with uncertainty quantification. Protein Sci. 31, e4467 (2022).

23. Lin, Z. et al. Evolutionary-scale prediction of atomic-level protein structure with a language model. Science 379, 1123–1130 (2023).

24. Dauparas, J. et al. Robust deep learning–based protein sequence design using ProteinMPNN. Science 378, 49–56 (2022).

25. Ferrandi, E. E. & Monti, D. Amine transaminases in chiral amines synthesis: recent advances and challenges. World J. Microbiol. Biotechnol. 34, 13 (2017).

26. Kelly, S. A., Mix, S., Moody, T. S. & Gilmore, B. F. Transaminases for industrial biocatalysis: novel enzyme discovery. Appl. Microbiol. Biotechnol. 104, 4781–4794 (2020).

27. Midelfort, K. S. et al. Redesigning and characterizing the substrate specificity and activity of Vibrio fluvialis aminotransferase for the synthesis of imagabalin. Protein Eng. Des. Sel. 26, 25–33 (2013).

28. Gao, S., Su, Y., Zhao, L., Li, G. & Zheng, G. Characterization of a (R)-selective amine transaminase from Fusarium oxysporum. Process Biochem. 63, 130–136 (2017).

29. Gomm, A. & O’Reilly, E. Transaminases for chiral amine synthesis. Curr. Opin. Chem. Biol. 43, 106–112 (2018).

30. Voss, M. et al. Creation of (R)-Amine Transaminase Activity within an α-Amino Acid Transaminase Scaffold. ACS Chem. Biol. 15, 416–424 (2020).

31. Telzerow, A. et al. Amine Transaminase from Exophiala Xenobiotica—Crystal Structure and Engineering of a Fold IV Transaminase that Naturally Converts Biaryl Ketones. ACS Catal. 9, 1140–1148 (2019).

32. Vaswani, A., et al. Attention Is All You Need. Preprint at 10.48550/arXiv.1706.03762 (2023).

33. Chai Discovery et al. Chai-1: Decoding the molecular interactions of life. Preprint at 10.1101/2024.10.10.615955 (2024).

34. Cui, Y. et al. Computational Redesign of a PETase for Plastic Biodegradation under Ambient Condition by the GRAPE Strategy. ACS Catal. 11, 1340–1350 (2021).

35. Cheng, P. et al. Zero-shot prediction of mutation effects with multimodal deep representation learning guides protein engineering. Cell Res. 34, 630–647 (2024).

36. Xu, Y., Liu, D. & Gong, H. Improving the prediction of protein stability changes upon mutations by geometric learning and a pre-training strategy. Nat. Comput. Sci. (2024) doi:10.1038/s43588-024-00716-2.

37. Chen, Y., Xu, Y., Liu, D., Xing, Y. & Gong, H. An end-to-end framework for the prediction of protein structure and fitness from single sequence. Nat. Commun. 15, 7400 (2024).

38. Jumper, J. et al. Highly accurate protein structure prediction with AlphaFold. Nature 596, 583–589 (2021).

39. Dieckhaus, H. & Kuhlman, B. Protein stability models fail to capture epistatic interactions of double point mutations. Protein Sci. 34, e70003 (2025).

40. Hopf, T. A. et al. Mutation effects predicted from sequence co-variation. Nat. Biotechnol. 35, 128–135 (2017).

41. Laimer, J., Hofer, H., Fritz, M., Wegenkittl, S. & Lackner, P. MAESTRO - multi agent stability prediction upon point mutations. BMC Bioinformatics 16, 116 (2015).

