## supplementary materials for "ProStab: Prediction of protein stability change upon mutations by protein language and inverse folding models"

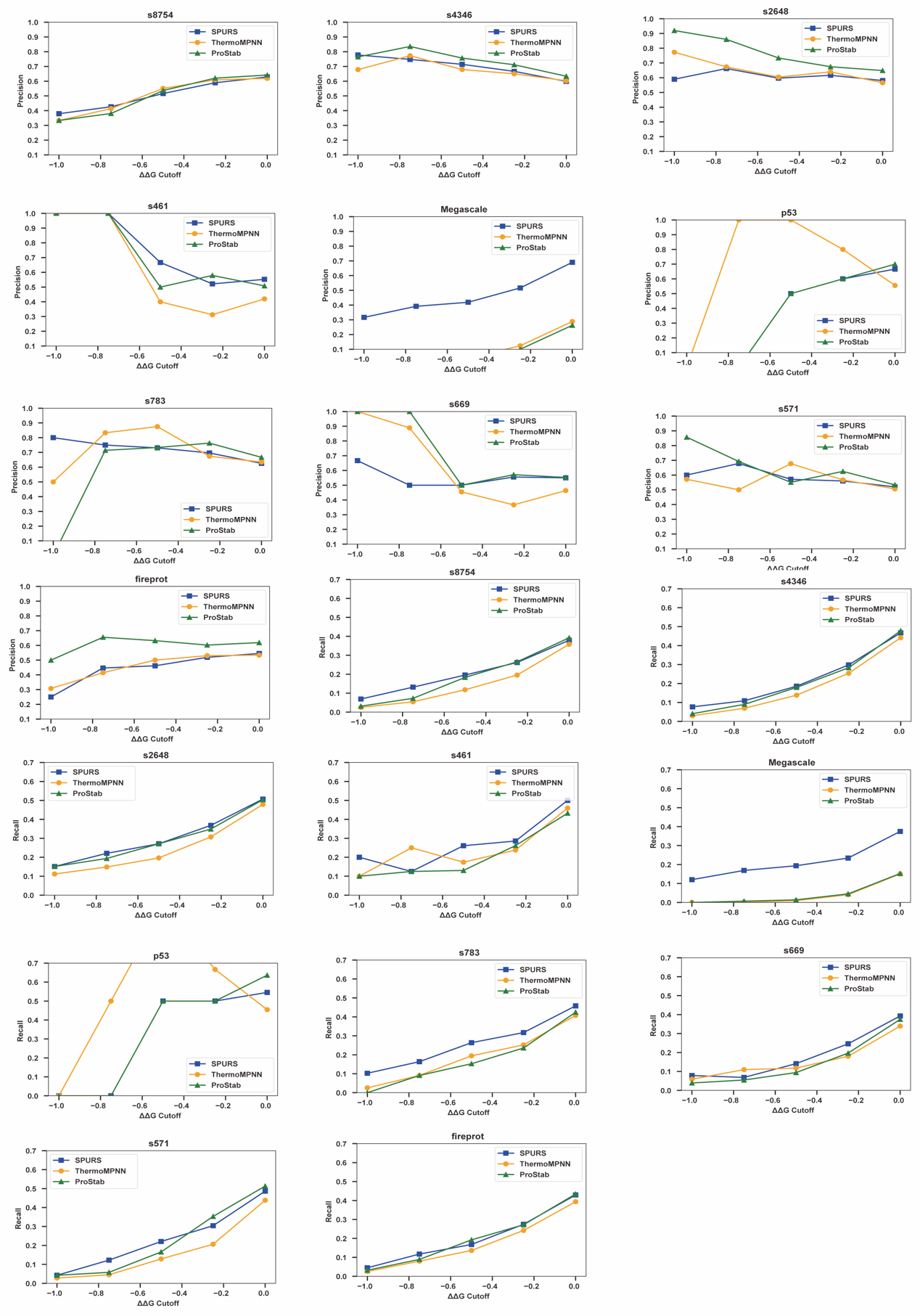

**Supplementary Figure 1. Benchmarking performance.** Precision and recall for ProStab, SPURS, and ThermoMPNN on ten test sets.

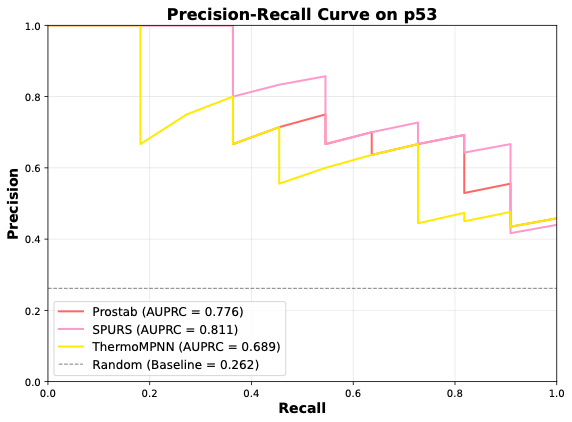

**Supplementary Figure 2.** Precision-recall curves for ProStab and two SOTA methods on P53 dataset.

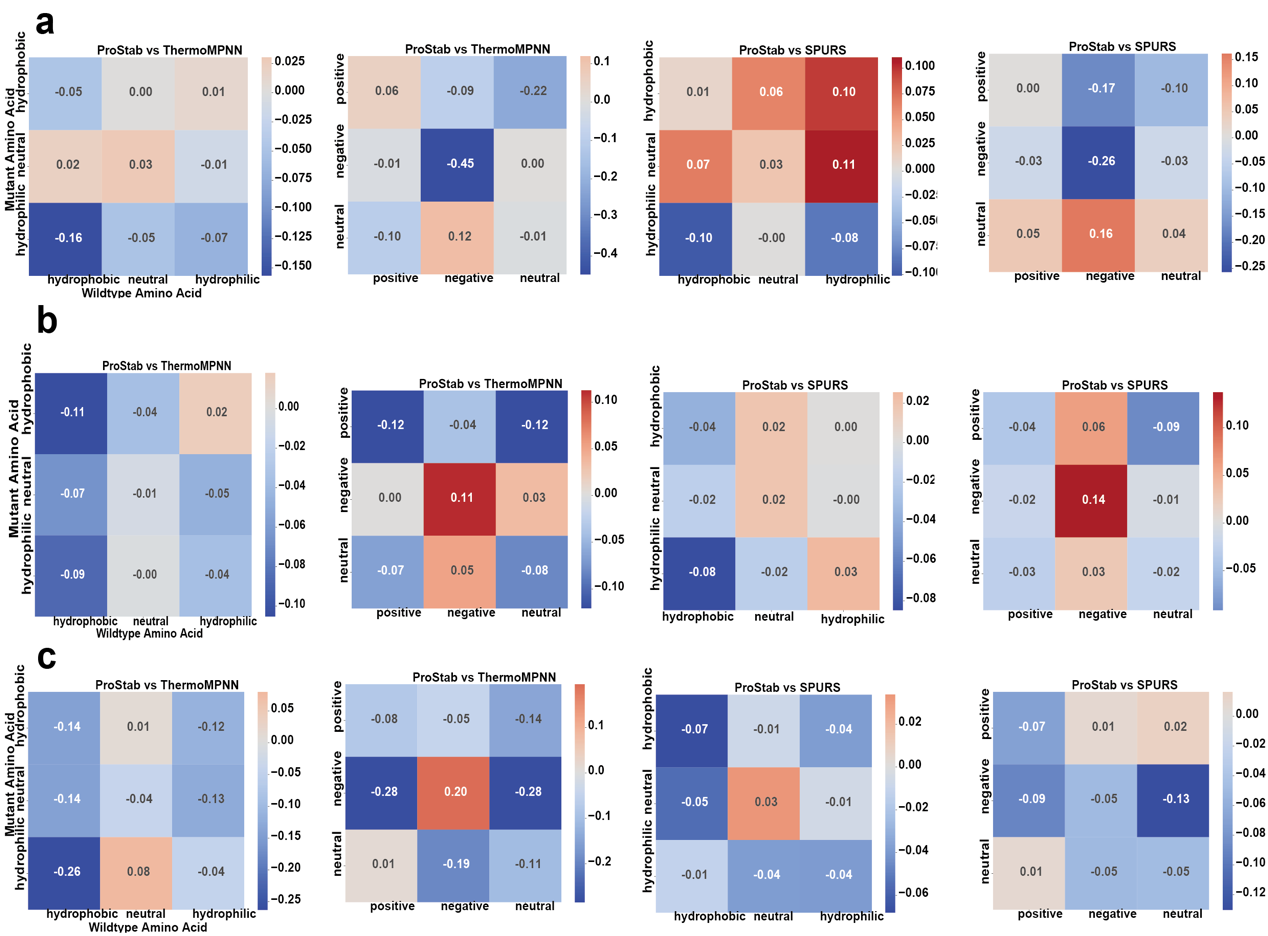

**Supplementary Figure 3. Benchmarking performance.** Precision and recall for ProStab, SPURS, and ThermoMPNN on ten test sets.

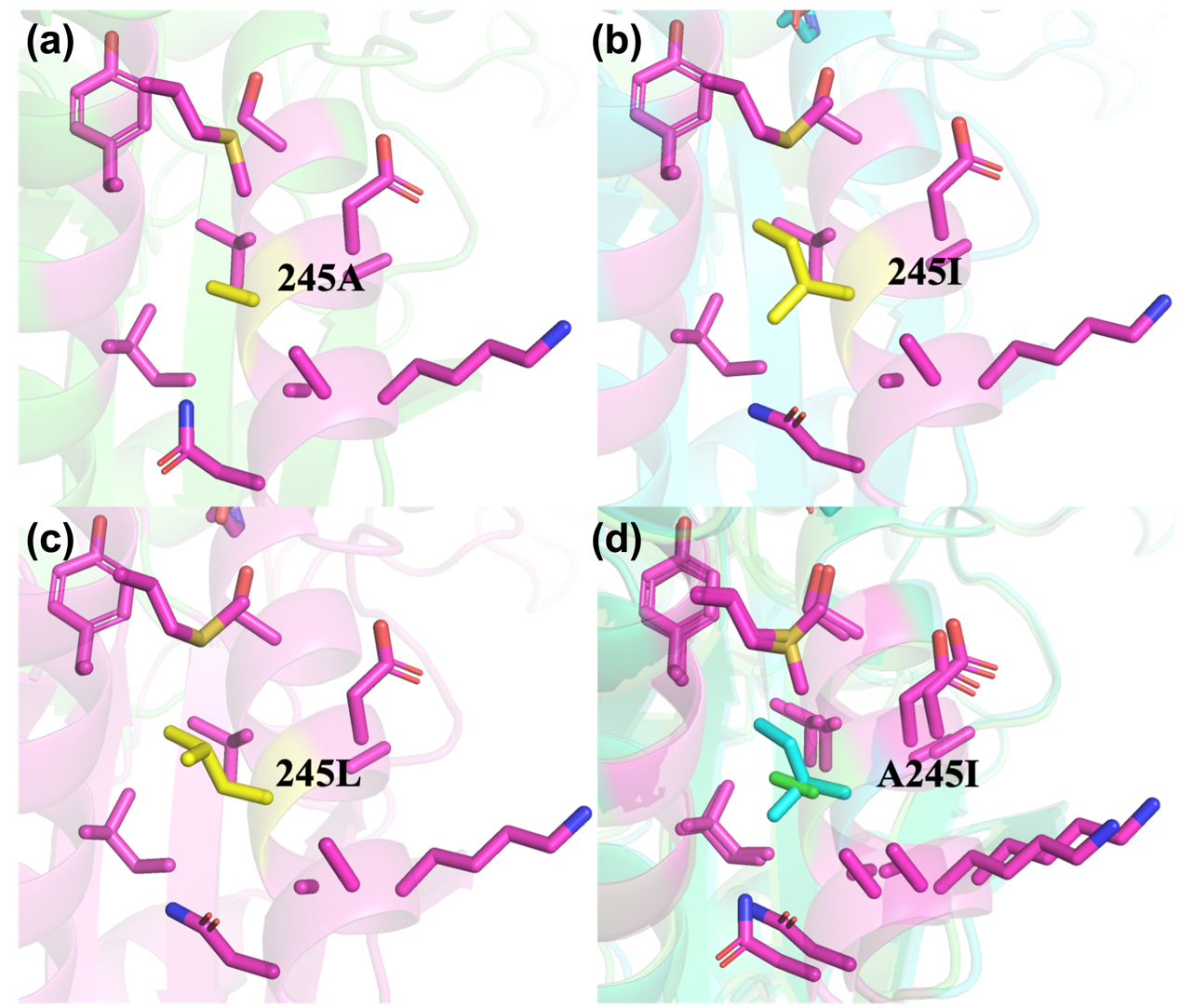

**Supplementary Figure 4. Comparison of WT and A245 mutants.** a. Predicted structure of the wild-type Ex-ATA and the arrangement of amino acid residues surrounding alanine at position 245. b. Predicted structure of the A245I mutant. c. Predicted structure of the A245L mutant. d. Structural comparison between the wild-type and the A245I mutant. The substitution of alanine at position 245 with leucine, which has a longer hydrophobic side chain, enhances hydrophobic interactions between the two alpha-helices, thereby improving protein stability.

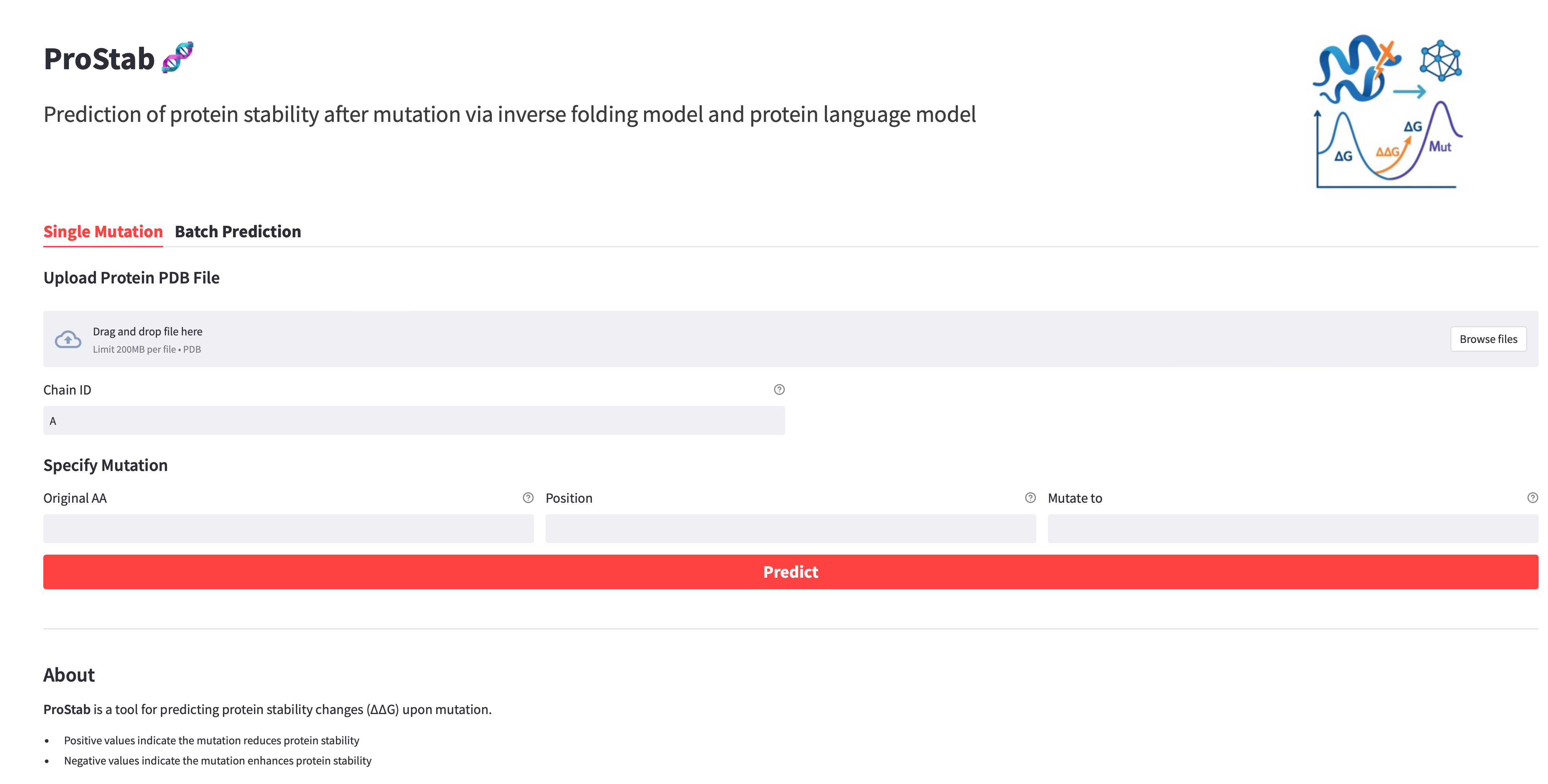

**Supplementary Figure 5. Overview of the online web server used for protein stability prediction.**

**Supplementary Table 1. Statistics of datasets used for model training and evaluation**. The number of unique proteins and total number of mutants across all proteins in each dataset.

| **Dataset** | **Number of proteins** | **Number of mutants** |
| --- | --- | --- |
| Megascale (training) | 212 | 203789 |
| Megascale (validation) | 31 | 27481 |
| Megascale (test) | 28 | 28312 |
| Fireprot (HF) | 89 | 3438 |
| S669 | 94 | 669 |
| S461 | 48 | 461 |
| S783 | 55 | 783 |
| S8754 | 274 | 8236 |
| S2648 | 131 | 2648 |
| S4346 | 281 | 3988 |
| S571 | 37 | 571 |

**Supplementary Table 2. The primers used in this study.**

| **Primer** | **Primer Sequence (5' to 3')** |
| --- | --- |
| N137-R | GTTGGTCAGGCTTTCGCC |
| N137R-F | GAAAGCCTGACCAACCGTCTGTATATGTGGATTCAGCCGT |
| E150-R | CATCACCCAAATATACGGCTGAATCC |
| E150P-F | TATATTTGGGTGATGCCGCCGGAAATGCAGCGCACC |
| P201-R | ATAATCTGCACCACGATCCTGC |
| P201V-F | CGTGGTGCAGATTATGTGTTTCTGACCGATGGCGACG |
| R239-R | AGTCACGCCTTTCAGCAC |
| R239L-F | CTGAAAGGCGTGACTCTGAAAAGCGTGGCAGATGCGGC |
| A243-R | CACGCTTTTGCGAGTCACG |
| A243I-F | ACTCGCAAAAGCGTGATTGATGCGGCGAAAGCGAAC |
| A245-R | ATCTGCCACGCTTTTGCGA |
| A245I-F | AAAAGCGTGGCAGATATTGCGAAAGCGAACGGCATT |
| A245L-F | AAAAGCGTGGCAGATCTGGCGAAAGCGAACGGCATT |
| A245V-F | AAAAGCGTGGCAGATGTGGCGAAAGCGAACGGCATT |
| N249-R | CGCTTTCGCCGCATCTG |
| N249L-F | GATGCGGCGAAAGCGCTGGGCATTGAAATGCGCATTGAAT |
| E300-R | TTTGGTCACCGGGCCCAC |
| E300L-F | GGCCCGGTGACCAAACTGATTTGGGATGGCTATTGGGC |
| E300M-F | GGCCCGGTGACCAAAATGATTTGGGATGGCTATTGGGC |
| G304-R | ATCCCAAATTTCTTTGGTCACCG |
| G304A-F | AAAGAAATTTGGGATGCGTATTGGGCGATGCATTATGACGA |
| G304K-F | AAAGAAATTTGGGATAAATATTGGGCGATGCATTATGACGA |
| G304L-F | AAAGAAATTTGGGATCTGTATTGGGCGATGCATTATGACGA |
| G304M-F | AAAGAAATTTGGGATATGTATTGGGCGATGCATTATGACGA |
| G304R-F | AAAGAAATTTGGGATCGTTATTGGGCGATGCATTATGACGA |
| G304W-F | AAAGAAATTTGGGATTGGTATTGGGCGATGCATTATGACGA |
| H309-R | CATCGCCCAATAGCCATCC |
| H309C-F | GGCTATTGGGCGATGTGCTATGACGATAAGTACAGCTT |
| H309I-F | GGCTATTGGGCGATGATTTATGACGATAAGTACAGCTT |
| H309V-F | GGCTATTGGGCGATGGTGTATGACGATAAGTACAGCTT |

**Supplementary Table 3. Plasmids used in this study.**

| **Plasmids** | **Relevant properties** | **Source** |
| --- | --- | --- |
| pET28a-*ExATA* | *ExATA* gene was synthesized and cloned to the NdeI and XhoI site of pET28a | Hongxun Biotech |
| pET28a-*ExATA*-N137R | Site-directed mutagenesis of asparagine 137 to arginine in *ExATA* | This study |
| pET28a-*ExATA*-E150P | Site-directed mutagenesis of Glutamic acid 150 to proline in *ExATA* | This study |
| pET28a-*ExATA*-P201V | Site-directed mutagenesis of proline 201 to valine in *ExATA* | This study |
| pET28a-*ExATA*-A243I | Site-directed mutagenesis of alanine 243 to isoleucine in *ExATA* | This study |
| pET28a-*ExATA*-A245I | Site-directed mutagenesis of alanine 245 to isoleucine in *ExATA* | This study |
| pET28a-*ExATA*-A245L | Site-directed mutagenesis of alanine 245 to leucine in *ExATA* | This study |
| pET28a-*ExATA*-A245V | Site-directed mutagenesis of alanine 245 to valine in *ExATA* | This study |
| pET28a-*ExATA*-N249L | Site-directed mutagenesis of asparagine 249 to leucine in *ExATA* | This study |
| pET28a-*ExATA*-G304A | Site-directed mutagenesis of glycine 304 to alanine in *ExATA* | This study |
| pET28a-*ExATA*-G304K | Site-directed mutagenesis of glycine 304 to lysine in *ExATA* | This study |
| pET28a-*ExATA*-G304L | Site-directed mutagenesis of glycine 304 to leucine in *ExATA* | This study |
| pET28a-*ExATA*-G304M | Site-directed mutagenesis of glycine 304 to methionine in *ExATA* | This study |
| pET28a-*ExATA*-G304R | Site-directed mutagenesis of glycine 304 to arginine in *ExATA* | This study |
| pET28a-*ExATA*-G304W | Site-directed mutagenesis of glycine 304 to tryptophan in *ExATA* | This study |
| pET28a-*ExATA*-H309C | Site-directed mutagenesis of histidine 309 to cysteine in *ExATA* | This study |
| pET28a-*ExATA*-H309I | Site-directed mutagenesis of histidine 309 to isoleucine in *ExATA* | This study |
| pET28a-*ExATA*-H309V | Site-directed mutagenesis of histidine 309 to valine in *ExATA* | This study |

**Supplementary Table 4. Specific activity and relative activity of wild-type and four mutant variants measured at various times.**

|  | **Time (min)** | **Concentration（mg/ml）** | **Specific Activity（U/mg）** | | | **Relative Activity** |
| --- | --- | --- | --- | --- | --- | --- |
| WT | 0 | 0.1 | 4.55 | 4.55 | 4.61 | 100.05% |
|  | 1 | 0.1 | 4.11 | 4.34 | 4.36 | 93.44% |
|  | 2 | 0.1 | 3.28 | 3.46 | 3.58 | 75.27% |
|  | 5 | 0.1 | 2.47 | 2.52 | 2.54 | 54.90% |
|  | 7 | 0.1 | 1.52 | 1.60 | 1.59 | 35.15% |
|  | 10 | 0.1 | 0.96 | 1.00 | 1.08 | 21.94% |
|  | 20 | 0.1 | 0.00 | 0.00 | 0.00 | 0.00% |
| P201V | 0 | 0.1 | 3.42 | 3.59 | 3.52 | 99.97% |
|  | 20 | 0.1 | 3.09 | 3.01 | 3.07 | 87.09% |
|  | 30 | 0.1 | 2.73 | 2.68 | 2.71 | 77.03% |
|  | 40 | 0.1 | 2.64 | 2.53 | 2.58 | 73.54% |
|  | 60 | 0.1 | 2.28 | 2.15 | 2.14 | 62.45% |
| N2499 | 0 | 0.1 | 4.25 | 4.31 | 4.27 | 99.98% |
|  | 20 | 0.1 | 3.20 | 3.32 | 3.22 | 75.90% |
|  | 30 | 0.1 | 2.69 | 2.82 | 2.96 | 65.99% |
|  | 40 | 0.1 | 2.73 | 2.78 | 2.86 | 65.19% |
|  | 60 | 0.1 | 2.34 | 2.35 | 2.41 | 55.30% |
| A245I | 0 | 0.1 | 3.19 | 3.19 | 3.26 | 99.88% |
|  | 20 | 0.1 | 3.02 | 2.95 | 2.92 | 92.08% |
|  | 30 | 0.1 | 2.96 | 2.90 | 2.91 | 90.73% |
|  | 40 | 0.1 | 2.91 | 2.81 | 2.85 | 88.75% |
|  | 60 | 0.1 | 2.94 | 2.81 | 2.81 | 88.64% |
| A245L | 0 | 0.1 | 3.48 | 3.44 | 3.62 | 100.05% |
|  | 20 | 0.1 | 3.05 | 3.09 | 3.09 | 87.65% |
|  | 30 | 0.1 | 2.90 | 2.96 | 2.98 | 83.90% |
|  | 40 | 0.1 | 2.80 | 2.88 | 2.86 | 81.08% |
|  | 60 | 0.1 | 2.69 | 2.78 | 2.75 | 78.03% |

**Supplementary Table 5. Enzyme activity of wild-type and mutant variants after incubation at 4°C for 5 minutes.** Specific activity was calculated from ΔA values, and the average of three replicates is shown.

| **Enzyme** | **Concentration (mg/mL)** | **ΔA (1)** | **ΔA (2)** | **ΔA (3)** | **Activity (1)** | **Activity (2)** | **Activity (3)** | **Mean Activity** |
| --- | --- | --- | --- | --- | --- | --- | --- | --- |
| WT | 0.1 | 1.44 | 1.443 | 1.478 | 3.892 | 3.9 | 3.995 | 3.929 |
| N137R | 0.1 | 0.382 | 0.391 | 0.393 | 1.032 | 1.057 | 1.062 | 1.05 |
| E150P | 0.1 | 1.819 | 1.857 | 1.874 | 4.916 | 5.019 | 5.065 | 5.0 |
| P201V | 0.1 | 1.265 | 1.329 | 1.301 | 3.419 | 3.592 | 3.516 | 3.509 |
| A243I | 0.1 | 1.321 | 1.318 | 1.33 | 3.57 | 3.562 | 3.595 | 3.576 |
| N249L | 0.1 | 1.574 | 1.595 | 1.581 | 4.254 | 4.311 | 4.273 | 4.279 |
| A245L | 0.1 | 1.287 | 1.272 | 1.339 | 3.478 | 3.438 | 3.619 | 3.512 |
| A245I | 0.1 | 1.18 | 1.182 | 1.208 | 3.189 | 3.195 | 3.265 | 3.216 |
| G304A | 0.1 | 1.798 | 1.773 | 1.824 | 4.859 | 4.792 | 4.93 | 4.86 |
| G304K | 0.1 | 1.239 | 1.253 | 1.288 | 3.349 | 3.386 | 3.481 | 3.405 |
| G304L | 0.1 | 1.165 | 1.188 | 1.195 | 3.149 | 3.211 | 3.23 | 3.196 |
| G304M | 0.1 | 1.872 | 1.894 | 1.953 | 5.059 | 5.119 | 5.278 | 5.152 |
| G304R | 0.1 | 0.947 | 0.962 | 0.973 | 2.559 | 2.6 | 2.63 | 2.596 |
| G304W | 0.1 | 1.084 | 1.034 | 1.098 | 2.93 | 2.795 | 2.968 | 2.897 |
| H309C | 0.1 | 0.327 | 0.327 | 0.337 | 0.884 | 0.884 | 0.911 | 0.893 |
| H309I | 0.1 | 0.231 | 0.23 | 0.229 | 0.624 | 0.622 | 0.619 | 0.622 |
| H309V | 0.1 | 0.272 | 0.268 | 0.273 | 0.735 | 0.724 | 0.738 | 0.732 |
| blank | 0.1 | 0.005 | 0.001 | 0.005 | 0.014 | 0.003 | 0.014 | 0.01 |

**Supplementary Table 6. Predicted ΔΔG values (in kcal/mol) for 20 point mutations.** Values are sorted and displayed in two rows for clarity.

| A245I | G304A | A245V | A245L | G304R | H309I | H309V | G304K | N137R | H309C |
| --- | --- | --- | --- | --- | --- | --- | --- | --- | --- |
| -1.437 | -1.203 | -1.103 | -1.099 | -1.094 | -1.088 | -1.029 | -0.998 | -0.996 | -0.962 |

| G304W | E300M | E150P | G304L | G304M | E300L | A243I | N249L | P201V | R239L |
| --- | --- | --- | --- | --- | --- | --- | --- | --- | --- |
| -0.956 | -0.955 | -0.944 | -0.933 | -0.930 | -0.922 | -0.913 | -0.900 | -0.892 | -0.867 |
